# Reduced MOV10 reveals novel functional cortical connections in an increased fear response

**DOI:** 10.64898/2026.05.18.725995

**Authors:** Temirlan Shilikbay, Aatiqa Nawaz, Menghan Sun, Megan Doon, Isabella Olmo, Leo Cumbie, Julius Benson, Baher Ibrahim, Nien-Pei Tsai, Daniel Llano, Jozien Goense, Howard Gritton, Stephanie Ceman

## Abstract

The RNA helicase MOV10 is highly expressed in developing brain, is present in synapses and is required for embryonic viability. A murine brain-specific knockout of MOV10 (*Mov10* Deletion) has a thickened cortex, abnormal dendritic arborization and enhanced fear memory. In human studies, *MOV10* is among the loci that is correlated with enhanced cortical brain volumes and is also significantly associated with substance dependence by epigenetic profiling. Here we demonstrate that *Mov10* Deletion mice show enhanced fear learning that is aligned with impaired structural connectivity of canonical fear circuits revealed by Diffusion Tensor Imaging. We propose a model where MOV10 loss leads to increased GABRA2 expression in the hippocampus and reduced anatomical connectivity that drives augmented fear learning. Memory reactivation is observable during fear memory retrieval as an overall increase in fMRI functional activity in cortical regions. Taken together, this framework identifies that enhanced fear in the MOV10 model is driven via a “corticalized” fear response during re-exposure to the training context that is not driven by the canonical fear circuit. These findings support a molecular basis for non-traditional enhanced learning mechanisms activated by fearful events that shed light on the intractability of fear memories with the potential to inform PTSD and substance dependence disorders.

## INTRODUCTION

MOV10 is an RNA-helicase known for its role in microRNA-mediated silencing, its anti-viral activity and as a suppressor of retrotransposition (1). MOV10 is required for the completion of gastrulation during embryogenesis; thus, a full body knockout animal is embryonic lethal (2). MOV10 expression is significantly elevated in postnatal brain (3). To determine its functional role in mediating behavior, in our previous work, we created a brain-specific knockout of the *Mov10* gene (*Mov10* Deletion) and observed thickening in cortical layers 2-4 and increased dendritic arborization of cultured hippocampal neurons (4). The *Mov10* Deletion mouse performed comparably to WT mice in a variety of behavioral paradigms with the notable exception of showing augmented fear memory. The *Mov10* Deletion mouse demonstrated increased memory in both cued and context fear conditioning tests. However, what was unknown were the physiological properties that support this enhancement, specifically the electrical properties of individual neurons, synaptic activity between neurons, and communication between different brain regions in the specific domain that could support enhanced fear memory. Fear memory responses depend on several factors related to proper synaptic functioning, the evidence of which comes from studies of gene knockout mice. For example, mice heterozygous for a voltage-gated sodium channel *Scn8a* has impaired intrinsic excitability and increased fear memory (5, 6). A similar phenotype is observed in mice with decreased expression of the potassium voltage-gated channel K_v_7.2 encoded by *Kcnq2* (6). Glutamate and gamma-aminobutyric acid (GABA) receptors are also implicated in fear memory, with metabotropic glutamate receptor 5 (mGluR5) and δ-subunit-containing GABA_A_ receptors (GABA_A_δR) knockout mice having impaired fear memory (7,8). Expression of these genes and others with similar functions was dysregulated in the absence of MOV10, suggesting that aberrant electrophysiology might be a potential connection between gene expression and *Mov10* Deletion increased fear memory (4). Finally, since MOV10 is a crucial cofactor for microRNA-mediated transcript expression (9,10), it is important to note that there is a large body of literature that demonstrates the role of miRNAs in fear memory. Inhibition of miR-29 enhanced trace fear memory stability (11) and induction of miR-144-3p rescued fear extinction in extinction impaired mice (12). In addition, conditional knockout (cKO) mice for *Dicer*, which is essential for miRNA biogenesis, has enhanced fear memory (13,14), consistent with the phenotype of the *Mov10* Deletion mice (4).These findings provide evidence that dysregulated cell signaling or transcript expression resulting in altered excitability can promote exaggerated fear responses (15–17)

Several regions in the brain have been strongly implicated in fear memory, namely the prefrontal cortex, hippocampus, amygdala, and periaqueductal gray area of midbrain. In this canonical fear network, the hippocampus and prefrontal cortex encode spatial and temporal information, the amygdala serves as the association hub of the aversive signal and the specific cue, which relays this information to the periaqueductal gray region to initiate freezing behavior (18–21). However, there is evidence of involvement of other regions including the striatum (22,23), anterior cingulate cortex (24), and cerebellum (25) suggesting other regions can be recruited to participate in mediating the expression of the fear response (19).

In this study, we explored the molecular underpinnings of the increased fear memory observed in the *Mov10* Deletion mouse from behavior to gene expression. We begin by investigating the rate of fear memory acquisition and extinction and then explore the structural and functional connectivity of the brain using magnetic resonance imaging. We then evaluate the intrinsic electrophysiological properties of the neurons using patch clamp, as well as their network-level activity using multielectrode arrays. Finally, we evaluate the transcriptomic changes at multiple timepoints in the development of the hippocampus, ultimately identifying increased expression of the *Gabra2* gene as the common feature present across development and is supportive of a role for GABAA in enhanced information flow and LTP in the hippocampus (26). Taken together, our results suggest that the *Mov10* Deletion mouse has a unique fear memory expression network mediated by enhanced functional connection between the cortical areas and midbrain, orchestrated by both MOV10-dependent structural changes and functional changes that are driven by an augmented inhibitory neuronal environment.

## METHODS

### Mouse husbandry

Mice were housed in standard IVC cages with water and food (Inotiv, Teklad, cat. #2918) available ad libitum. Mice were kept on a 12-h light/dark cycle from 7 AM to 7□PM and 7□PM to 7□AM, respectively. All experiments involving mice were reviewed and approved by the University of Illinois at Urbana-Champaign Institutional Animal Care and Use Committee based on the recommendations of the Guide for the Care and Use of Laboratory Animals of the National Institutes of Health. Credentials of the IACUC committee include USDA Registration: #33-R-0029; PHS Assurance: D16-00075 (A3118-01); AAALAC: #00766. Experiments were approved in IACUC protocols 19112, 22113, and 25082 (07/28/2022–5/14/2028). Mice were euthanized by carbon dioxide asphyxiation by replacing air in their home cage with CO_2_ at the rate of 70% volume/min until they ceased moving and stopped breathing. Death was confirmed by cervical dislocation. For neuron dissection, P0 pups were euthanized by decapitation.

### Fear conditioning tests

Mice aged 8–12 weeks old of both sexes were tested. A modified procedure of the test was performed as described (4). Mice were trained by exposing them for 5 min to a chamber (34□×□28□×□30 cm) where they received consecutive foot shocks (0.5 mA, 2 s) with a 3-min break between each shock. The number of shocks varied between tests. For fear acquisition test, mice received five foot shocks while their behavior was recorded during the training. For fear extinction test, mice received three foot shocks, and their behavior in the original training chamber was recorded every day for four days after training. For context fear conditioning test as a part of MRI procedure, mice received five foot shocks, and their behavior in the original training chamber was recorded one day after the training right before the scanning. For long-term memory test, mice received five foot shocks, and their behavior in the original training chamber was recorded 30 days after training. Freezing was defined as lack of movement for at least 1 second, except for respiration. Percent of time spent freezing was used for the analysis. All trials for each mouse were videotaped with a Logitech HD Pro webcam and analyzed in TopScan, Cleversys Inc. software.

### Hot plate test

Mice aged 8–12 weeks old of both sexes were tested. A modified procedure of the test was performed as described (27). Mice were placed on a hot plate preheated to 55°C until the first nociceptive reaction which was licking of the hind paw. Every trial was recorded using a Logitech HD Pro webcam and the time between the start of the test and the nociceptive reaction onset was quantified manually.

### RNA sequencing and analysis

Three samples for both WT and *Mov10* Deletion were prepared by dissecting hippocampi of at least two P0 pups per sample and cryopreserving them in liquid nitrogen. The cryo-preserved sampled were shipped in dry ice to Biostate.ai. There, the RNAs were isolated and enriched for mRNAs using NEBNext Poly(A) mRNA Magnetic Isolation Module. RNA-sequencing libraries were prepared using Illumina kit and 150 bp reads were generated using Illumina NovaSeq 6000. After adapter using cutadapt (v. 3.7) -a AGATCGGAAGAGCACACGTCTGAACTCCAGTCA -AAGATCGGAAGAGCGTCGTGTAGGGAAAGAGTGT and quality checking using multiQC (v1.28), abundance of each transcript was quantified using the Selective Alignment method of Salmon (v1.10.0) with a decoy-aware transcriptome using the entire GRCm39 genome as the decoy. The differential expression analysis was performed in R (v. 4.5.1) using DESeq2 (1.48.1). Only genes with at least 10 counts in at least 3 samples were considered for the analysis, and the data was modeled by sample condition which is a single metric describing sample’s genotype and timepoint. Multiple testing correction was done using the False Discovery Rate method. Gene ontology analysis was performed using PANTHER (v. 19.0).

### Western blotting

Hippocampi from at least two P2 WT and *Mov10* Deletion per pups per sample were dissected, lysed in lysis buffer (50 mM Tris–Cl 7.5, 300 mM NaCl, 30 mM ethylenediaminetetraacetic acid (EDTA), 0.5% Triton), quantified by Bradford assay, and resuspended in 1□×□sample buffer for resolution by SDS-PAGE (7.5% gels) and analyzed by immunoblotting. Briefly, membranes were blocked with 5% non-fat dry milk in phosphate-buffered saline (PBS) containing 1% TWEEN-20 for 1 h at room temperature. Primary antibody was applied overnight at 4 °C followed by a brief wash in 1% non-fat milk PBS containing 1% TWEEN-20 wash buffer. Horseradish peroxidase (HRP)-conjugated secondary antibody was applied at 1:5000 dilution for 1 h at room temperature and washed 4□×□15 min using wash buffer. The HRP signal was detected using an enhanced chemiluminescent (ECL) substrate on BioRad ChemiDoc. The primary antibodies used were anti-MOV10 (RRID:AB_1040002, A301-571A, Bethyl Laboratories, Montgomery, TX, USA) at 1:1000, anti-GAPDH (AB_3670783, ab9484, Abcam) at 1:1000, anti-GABRA2 (RRID:AB_1040002, 83057-2-RR, Proteintech, Rosemont, IL, USA) at 1:2500, and HRP-conjugated goat anti-rabbit (RRID:AB_2337937, 111–035-008, Jackson Immunoresearch) and goat anti-mouse (RRID:AB_2338512, 115–035–174, Jackson Immunoresearch). The density of the bands was quantified using ImageJ. The level of significance and tests performed are described in the figure legends for each experiment.

### Whole cell patch clamp

The whole-cell patch clamp recordings on the hippocampal neurons were performed as previously described (28). Hippocampi of P0 pups were dissected and dissociated as described. Coverslips were coated for 2 h at room temperature with 10 μg/mL of poly-L-lysine (P4707, Sigma) and 330 cells/mm^2^ were plated in minimum essential medium (MEM) supplemented with 10% fetal bovine serum (FBS). After 24 h, the medium was switched to maintenance medium consisting of Neurobasal medium (21103049, Gibco) supplemented with B-27 (17504044, Gibco). Half of the media was removed and replaced with fresh NB medium every 3-4 days until DIV13-14. Then, the coverslip was transferred to the recording chamber in external solution containing (in mM): 126 NaCl, 26 NaHCO_3_, 2.5 KCl, 1.25 NaH_2_PO_4_, 2 MgCl_2_, 2 CaCl_2_, and 10 Dextrose, bubbled with 95% O_2_ and 5% CO_2_ (300–310 mOsm). The hippocampal neurons were visually identified using an upright microscope (Olympus BX51WI). The whole-cell patch-clamp recordings were carried out at room temperature in external solution containing the synaptic transmission blockers CNQX (5 μM), DL-AP5 (25 μM) and bicuculline (5 μM) for action potential (AP) under current clamp mode. Recording pipettes were pulled from borosilicate glass capillaries (King Precision Glass, Inc, glass type 7740, OD 1.5 mm) with an outer diameter of 1.5 mm on micropipette puller (P-97; Sutter Instruments, Novata, CA, USA), and had a resistance of 6-8 MΩ when filled with internal solution containing (in mM): 117 potassium gluconate, 1 MgCl_2_, 0.07 CaCl2, 0.1 EGTA, 13 KCl, 10 HEPES, 0.4 ATP, and 0.4 GTP. (290 mOsm, pH 7.3). Resting membrane potential was measured in current-clamp after achieving the whole-cell configuration. The action potential (AP) firing rates were measured upon delivering constant current pulses of 500 ms ranging from 0 to 200 pA in 20 pA increments with a step interval of 2 sec. Whole-cell recordings were made using a Multiclamp 700B amplifier (Molecular Devices). Electrophysiological recordings were filtered at low-pass Bessel at 10 kHz and digitized at 10 kHz. Data was acquired and analyzed with a Digidata 1550B interface (Molecular Devices) and the pClamp suite of software (version 11.4; Molecular Devices). Recording analyses were performed using Clampfit software (version 11.4; Molecular Devices) or pyabf (v. 2.3.8) in Python (v. 3.12.4). The instantaneous AP firing rate at each current pulse was calculated as a reciprocal of the interspike intervals (ISI), measured as the time between the first and second AP peaks (i.e. 1/interspike interval). The AP threshold was measured as the membrane potential at which dV/dt of the AP exceeded 10 mV/ms. The AP amplitude was calculated as the difference between the AP peak amplitude and the AP threshold. The 10–90% rise time and decay time were determined by calculating the difference between the time of 10% and 90% of the AP peak amplitude, respectively. The AP half width was measured as the width at half-maximal AP amplitude. Fast afterhyperpolarization (fAHP) was measured from trains of APs elicited by 100-pA current injection as the difference between the AP threshold and minimum voltage after the AP peak. The AP latency was assessed as the time between onset of current injection to peak of the first ensuing AP. Rheobase current was quantified as the minimal depolarizing current step sufficient to elicit an AP as described. Input resistance of the neuron was measured by dividing the average voltage deflection of the membrane potential by a hyperpolarizing current.

### MEA recording

Multielectrode array recordings were performed using Maestro Edge (Axion Biosystems) with Cytoview-MEA-6 plates (6-well plates) as described previously (29). Hippocampi from P0 mice were dissected, dissociated into single-cells, and cultured on MEA plates at 100,000 cells/well density in minimum essential medium (MEM) supplemented with 10% fetal bovine serum (FBS). After 24 h, the medium was switched to maintenance medium consisting of Neurobasal medium (21103049, Gibco) supplemented with B-27 (17504044, Gibco). Half of the media was removed and replaced with fresh maintenance medium every 3-4 days until DIV13-14. On DIV13-14, field potentials were recorded at each electrode relative to the ground electrode with a sampling frequency of 12.5 kHz. Followed by 30 min baseline recording of hippocampal cultures without any treatment, neurons were treated with Bicuculline (20 μM), Muscimol (75 nM), DHPG (100 μM), MPEP (10 μM), and LY367385 (100 μM) as indicated and immediately recorded for another 30 min. To avoid the effect physical disturbance of the culture on the network activity, only the last 15 min of the recordings were used for data analysis. Axis Navigator software (v. 3.4.1.15, Axion Biosystems) was used for spike extraction from raw electrical signals. Spike detector setting for each electrode was independently set at the threshold of ±6 standard deviation. Therefore, activity above the threshold was counted as a spike and included in data for analysis. Data from MEA wells with the number of active electrodes (5 spikes/minute) less than 15 out of 64 were omitted from the analysis. For burst detection, a minimum of 5 spikes with a maximum 100 ms interspike interval was set for individual electrodes.

### MRI acquisition and analysis

MRI scans were performed using a 9.4 Tesla preclinical MRI system (BioSpec 94/30 USR, Bruker BioSpin MRI, Billerica, Massachusetts, USA) equipped with a B-GA12S HP gradient insert optimized for rodent imaging, a 12-cm volume transmit RF-coil and a mouse brain 2x2 phased array receive coil. Prior to the experiment, mice were anesthetized using 4-5% isoflurane in oxygen inside an induction chamber using an isoflurane vaporizer (EZ-SA800, E-Z Systems Inc., Palmer, Pennsylvania, USA). Mice were placed in the mouse bed equipped with a bite bar and ear bars for head fixation. Surgical silicone adhesive (Low Toxicity Silicone Adhesive, World Precision Instruments, cat. KWIK-SIL) with petroleum jelly were put on the shaved mouse head to reduce image distortions. Mice received an IP injection of medetomidine (1 mg/kg). Anesthesia was supplemented with 0.25-0.5% isoflurane. Body temperature was maintained within nominal range using a circulating water-based heating system. Body temperature and respiratory function were tracked continuously using an MRI-compatible animal monitoring & gating System (Model 1030, SA Instruments. Inc., Stony Brook, New York, USA). T_2_-weighted scans were acquired using a RARE sequence with 3200 ms TR, 44 ms TE, 256x256x30 matrix, 75x75x500 μm^3^ resolution, 90° flip angle for a total of 7.5 min. Resting state fMRI scans were acquired using a single-shot gradient-echo EPI sequence with 1500 ms TR, 14 ms TE, 64x64x30 matrix, 300x300x500 μm^3^ resolution, 65° flip angle for a total of 12.5 min. DTI scans were acquired using a 4-segment DTI-EPI sequence with 3000 ms TR, 32 ms TE, 64x64x30 matrix, 150x150x500 μm^3^ resolution, 90° flip angle, 60 directions, 1000 s/mm^2^ and 2000 s/mm^2^ b-factors for a total of 25 min. The MRI data were processed using a custom-built pipeline (https://github.com/temshil/mri) based on AIDAmri (30). fMRI data was preprocessed using FSL MCFLIRT for motion correction, FSL TOPUP for phase-encoding distortion-correction, FSL MELODIC for regressing out physiological noise (31). DTI data was preprocessed using Patch2Self (32) to remove thermal noise, FSL EDDY to remove distortions, and FSL TOPUP for phase-encoding distortion-correction. The Allen Brain Atlas (33) with custom parcellations was registered to the fMRI and DTI data for connectivity analysis using NiftyReg (34). DTI data was processed in DSI Studio (35) using previously published parameters optimized for mouse brains (36) to generate tracts. Functional and structural correlations, graph analysis and statistical analysis were conducted in Python (v. 3.12.4) using nibabel (v. 5.2.1), bctpy (v. 0.6.1), scipy (v. 1.14.1), respectively. Circos plots were plotted using pycirclize (v. 1.10.1). For DTI analysis, since we did not anticipate the behavioral test to change the morphology of the brain, we used the 10 scans of the highest quality, regardless of whether the scan was performed before or after fear conditioning. For fMRI analysis, following rigorous preprocessing, three datasets were excluded due to persistent motion and physiological artifacts that could not be sufficiently removed. Therefore, subsequent analyses were conducted on a final sample of 11 out of 14 scans.

Structural connectivity (SOC) was measured using the following formula to quantify the structural connectivity between the different brain regions: 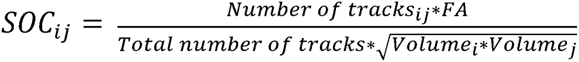. Briefly, SOC represents the number of tracts terminating between two brain regions weighted to their FA values to account for tracts’ integrity and normalized to the total number of generated tracts to account for scan-specific differences and the geometric mean of the two brain regions to account for size differences of the connected brain regions.

### Statistical analysis

Data for each experiment were obtained independently through random sampling. Prior to statistical analysis, normality was assessed using the Shapiro–Wilk test. In cases where the assumption of normality was violated, the Mann–Whitney U test was employed for comparisons involving two groups. For data that met the normality assumption but exhibited unequal variances, Welch’s t-test was utilized for two-group comparisons. When both normality and homogeneity of variance were satisfied, Student’s t-test was used for two-group comparisons. For comparing the slopes of the behavioral data trendlines, we calculated standard error of the slopes as *s(b)* = 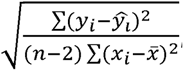, where *n* is the total sample size, *y_i_* is the observed dependent variable, 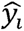 is the predicted dependent variable, *x_i_* is the independent variable, 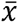 is the mean value of the independent variables. The degree of freedom was estimated using Satterthwaite approximation. For MRI analysis, none of the FDR-adjusted p-values were significant, a common concern in the field (37,38). Thus, we plotted the significantly changed region pairs using raw p-values for both DTI and fMRI results. All statistical analyses were conducted in R (v4.5.1) or Python (v. 3.12.4).

## RESULTS

To investigate the dynamics of fear memory in the *Mov10* Deletion mouse, we first looked at the rate of fear learning in a fear conditioning paradigm (Fig. 1A). To compare the rate of learning, we used a logarithmic fit and found that the *Mov10* Deletion mice displayed a significantly higher rate of fear learning than WT mice as indicated by a larger fit coefficient (Fig. 1A: 35.91 vs 22.76). Larger trendline coefficients indicate that the *Mov10* Deletion mice form new memories faster than the WT mice. Similarly, the *Mov10* Deletion mice showed a more rapid decline in extinction based on their substantially higher freezing rate on the first day (Fig. 1B, -19.36 vs -15.11), that continued in all the fear extinction trials, indicating that the fear memory formed during the conditioning training persisted significantly longer in the *Mov10* Deletion mice.

**Figure 1.**
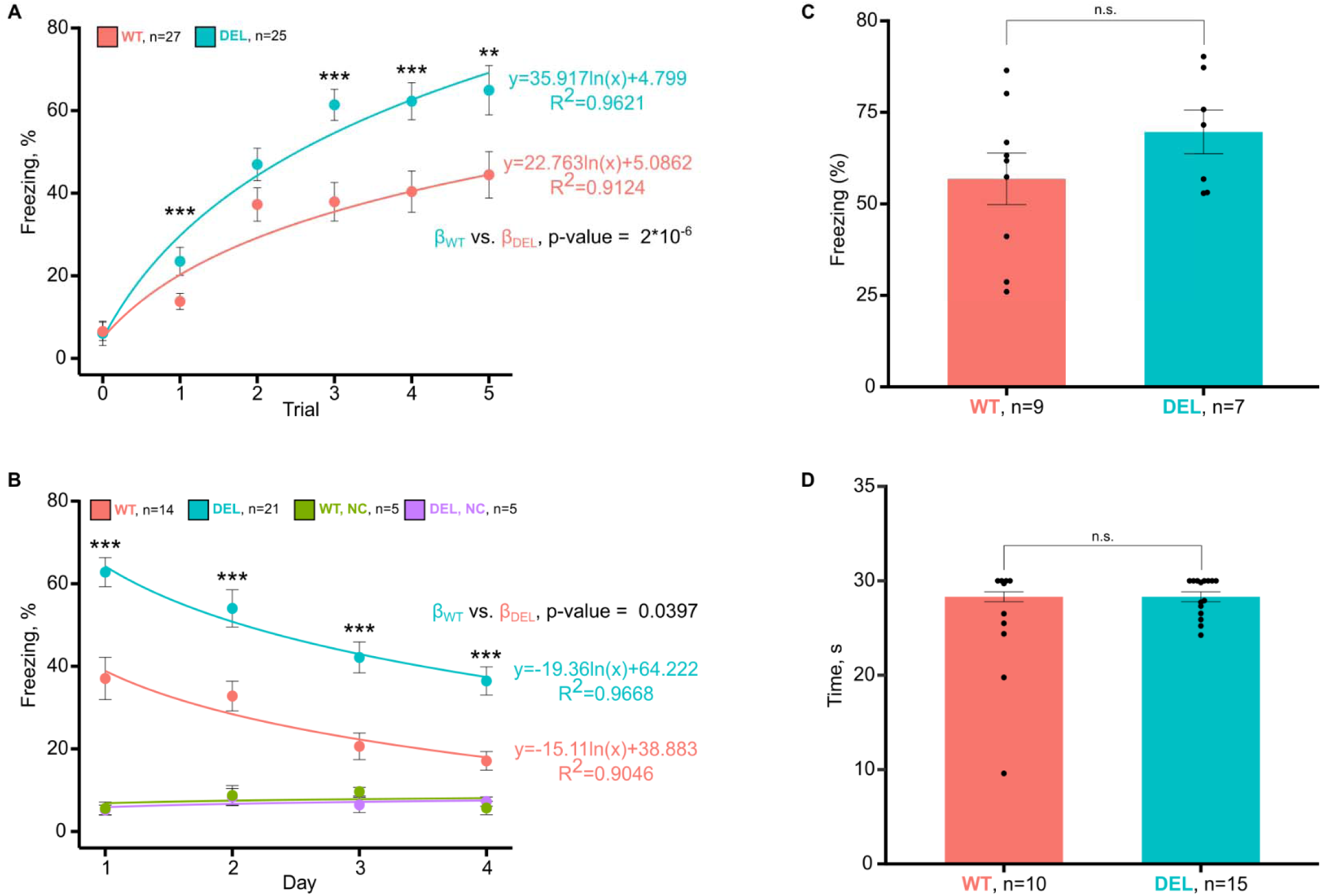
*Mov10* Deletion mice have increased fear learning and extinction. **A)** Fear memory acquisition test. Y axis is % of time spent freezing and the X axis is the trial number representing the time between consecutive shocks (3 minutes). **B)** Fear extinction test. Y axis is % of time spent freezing and X axis is the number of days after fear training. **C)** Long term fear memory test. Y axis is % of time spent freezing 30 days after fear training. X axis indicates genotype. **D)** Nociception test. Y axis is time in seconds (s) before responding to heat (hindpaw licking). X axis indicates genotype. *Mov10* Deletion (DEL). Data are shown as mean□±□SEM. **p-value<0.01, ***p-value<0.001, n is the number of mice of the genotype indicated, both sexes from *N*□>□3 litters.

To test if the *Mov10* Deletion mice have increased fear memory retention compared to WT mice, we performed a long-term memory test 30 days later. Although the *Mov10* Deletion mice displayed a higher average percent freezing than WT, the difference was not significant, indicating that memory decay in *Mov10* Deletion mice remained elevated in both WT and *Mov10* Deletion mice (Fig. 1C). Finally, to determine if the increased fear memory was caused by increased sensitivity to the pain of foot shock (39), we performed a classic nociception test (27) and found no difference between genotypes (Fig. 1D). Thus, the enhanced fear learning observed in the *Mov10* Deletion mouse must be due to differences in signal processing in the brain.

We then employed magnetic resonance imaging (MRI) to examine brain connectivity in an unbiased manner in a live mouse by scanning mice of both genotypes using T_2_*-weighted, resting-state functional MRI (rs-fMRI), and diffusion tensor imaging (DTI) sequences twice—once on the naïve mice and once 24 hours later and immediately after a 15 minute re-exposure to the training context (Fig. 2A). We first compared the structural connectivity of the *Mov10* Deletion and WT mice and did not observe any differences in the fractional anisotropy (FA), mean diffusivity, axial diffusivity, and radial diffusivity across any brain regions (Supplementary Fig. 1A-D). Differential tractography analysis did not identify any tracts with significantly different quantitative anisotropy, and we did not see any difference in the graph analysis measures (Supplementary Fig. 2E). To compare the number of tracts connecting all the brain regions, we calculated the strength of connectivity (SOC), described in the Methods. The average and standard deviation matrices are similar between WT and *Mov10* Deletion mice (Fig. 2B, Supplementary Fig. 2A-C), indicating high consistency across scans both within and across the two genotypes. However, analysis of the SOC at each region pair found reduced connections in the *Mov10* Deletion mice in the canonical fear network regions (Fig. 2C). Representative tracts connecting the amygdala and hippocampus in particular within the same hemispheres were reduced in *Mov10* Deletion mice explaining the reduced across-node structural connectivity (Fig. 2D). We also identified reduced average connectivity in cortical regions (Supplementary Fig. 2D) in the *Mov10* Deletion mice. In summary, our survey of the structural connectivity in the *Mov10* Deletion brain suggests that multiple regions, specifically, the cortical areas and the canonical fear network regions, have reduced structural connectivity compared to the WT brain.

**Figure 2.**
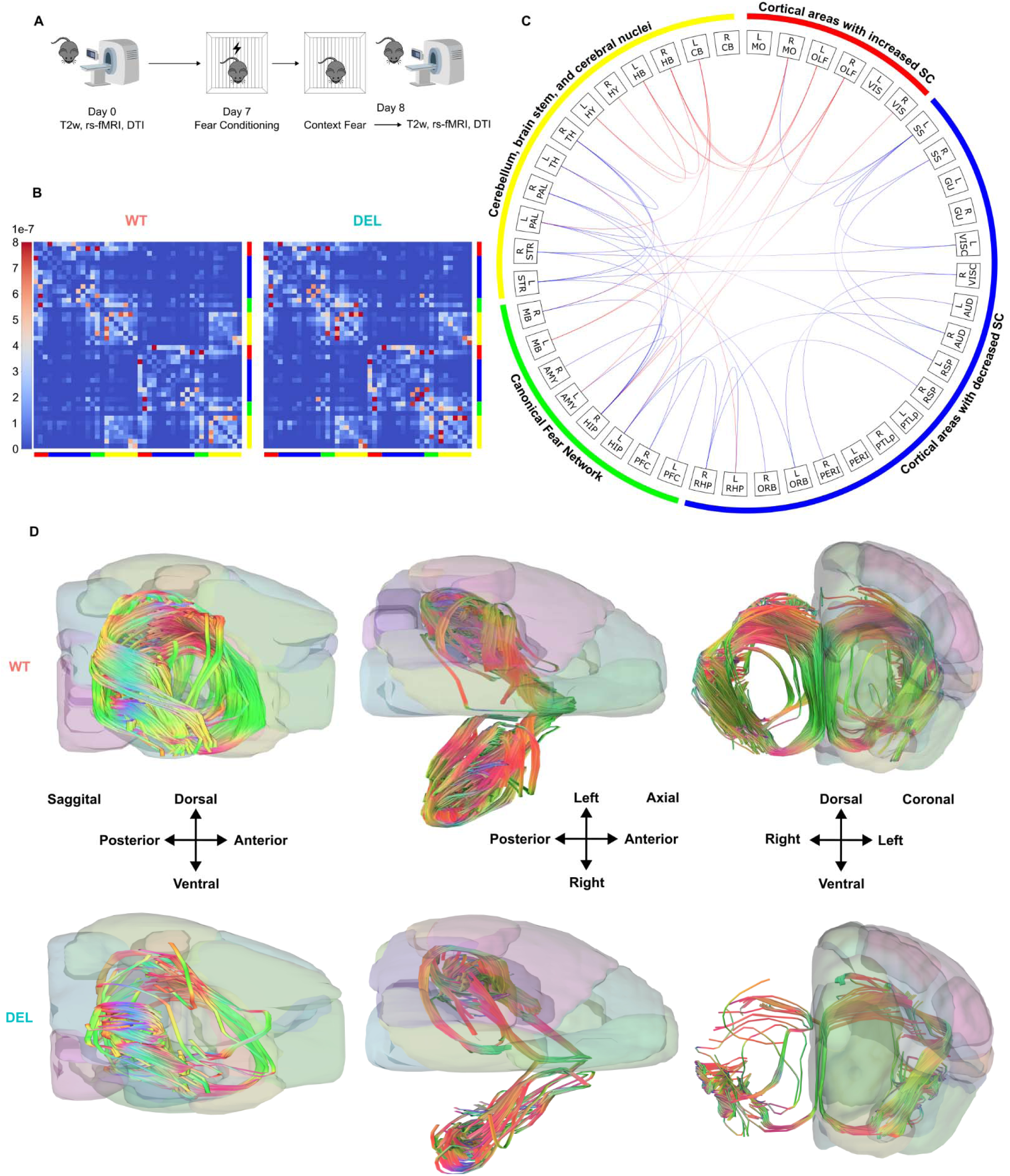
*Mov10* Deletion mice have reduced structural connectivity between multiple brain regions. **A)** Schematic of the MRI timeline. The first scan was performed on day 0 and the second was performed eight days later. **B)** Average connectivity matrices of strength of connectivity (SOC) between 22 bilateral brain regions across 10 animals for each genotype. *Mov10* Deletion (DEL). DTI data were preprocessed using a custom in-house script, registered to the modified Allen Brain Atlas containing 22 bilateral regions (Supplementary Table 1), and analyzed in DSI Studio (35). **C)** Circos plot showing the significantly (p-value<0.05) reduced (blue) or significantly increased (red) structural SOC in *Mov10* Deletion (DEL) compared to WT. R-right; L-left. Cortices: MO-Motor; SS-Somatosensory; GU-Gustatory; VISC-Visceral; OLF-Olfactory; AUD-Auditory; VIS-visual; RSP-Restrosplenial; PTL-Posterior Parietal; PERI-Perirhinal; PFC-Prefrontal; ORB-orbital; HIP-hippocampus; RHP-RetroHippocampal region; STR-Striatum; PAL-Pallidum; TH-Thalamus; HY-Hypothalamus; AMY-Amygdala; MB-Midbrain; HB-Hindbrain; CB-Cerebellum; SC-Structural Connectivity. **D)** Representative tracts passing between R AMY and R HIP and between L AMY and L HIP in WT (top) and *Mov10* Deletion (DEL) (bottom). Brain regions of the left hemisphere are shown in a semitransparent opacity for reference.

We then examined the rs-fMRI data acquired from naïve mice of both genotypes. Overall, the average and standard deviation matrices of Pearson correlation coefficients were highly consistent across genotypes, with high connectivity between the same contralateral regions and within the different regions on the same hemisphere (Fig. 3A, Supplementary Fig. 3A-B). However, closer analysis revealed multiple brain region pairs had both increased and decreased functional connectivity in the naïve mice (Fig. 3B), the pairs of regions with significant changes did not show any notable enrichment, and averaging the connectivity coefficients mitigated the differences (Supplementary Fig. 3C). Thus, we concluded that the brain activity of naïve *Mov10* Deletion and WT mice on average was highly similar. Our previous work showing that the *Mov10* Deletion mice did not have increased hyperactivity, anxiety, or spatial and object memory supports this conclusion (4).

**Figure 3.**
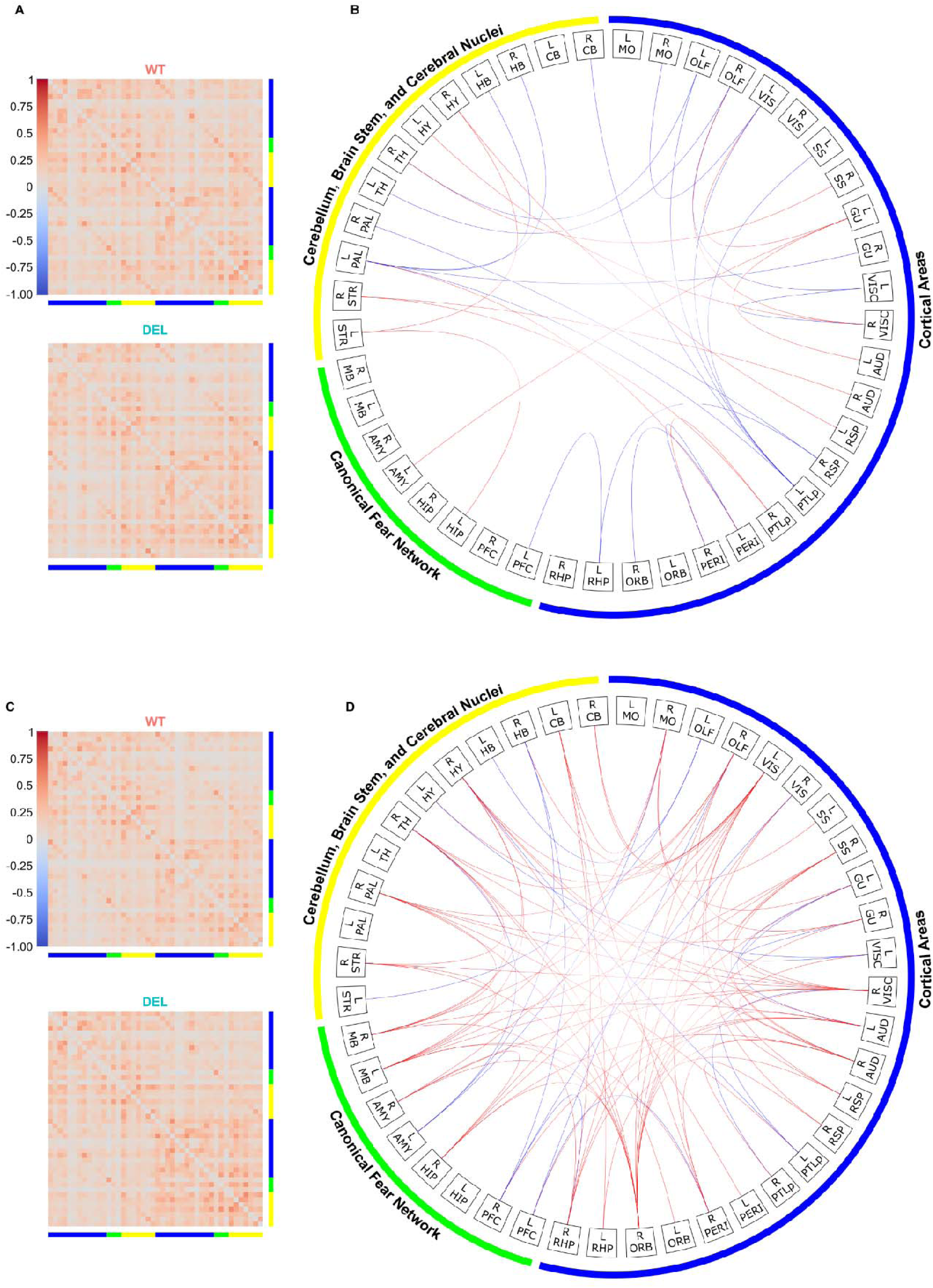
*Mov10* Deletion mice have increased functional connectivity between multiple brain regions after re-exposure to context. **A)** Average functional connectivity matrices of Pearson correlation coefficient between 22 bilateral brain regions across 11 animals for each genotype calculated from fMRI data acquired before fear conditioning. *Mov10* Deletion (DEL). **B)** Circos plot showing the significantly (p-value<0.05) reduced (blue) or significantly increased (red) functional correlation in *Mov10* Deletion compared to WT calculated from fMRI data acquired before fear conditioning. R-right; L-left. Cortices: MO-Motor; SS-Somatosensory; GU-Gustatory; VISC-Visceral; OLF-Olfactory; AUD-Auditory; VIS-visual; RSP-Restrosplenial; PTL-Posterior Parietal; PERI-Perirhinal; PFC-Prefrontal; ORB-orbital. HIP-hippocampus; RHP-RetroHippocampal region; STR-Striatum; PAL-Pallidum; TH-Thalamus; HY-Hypothalamus; AMY-Amygdala; MB-Midbrain; HB-Hindbrain; CB-Cerebellum. **C)** Average functional connectivity matrices of Pearson correlation coefficient between 22 bilateral brain regions across 11 animals for each genotype calculated from fMRI data acquired after re-exposure to context. *Mov10* Deletion (DEL). **D)** Circos plot showing the significantly (p-value<0.05) reduced (blue) or significantly increased (red) functional correlation in *Mov10* Deletion compared to WT calculated from fMRI data acquired after re-exposure to context.

Next, we examined how the WT and *Mov10* Deletion mice responded to re-exposure to the training chamber 24 hours after contextual fear conditioning, i.e., during memory retrieval (Supplementary Fig. 3D). Graph analysis performed on the rs-fMRI data did not find any significant differences between genotypes or before and after fear conditioning (Supplementary Fig. 4A-B). The average and standard deviation matrices of Pearson correlation coefficients from WT and *Mov10* Deletion data showed the same pattern as for the rs-fMRI data collected before the fear conditioning test, suggesting high consistency across scans (Fig. 3C, Supplementary Fig. 3E-F). However, the non-FDR-corrected p-values showed that the majority of brain regions in the *Mov10* Deletion mice had increased functional connectivity compared to the WT mice (Fig. 3D). These included both cortical regions (somatosensory, visual, retrosplenial, auditory, orbital) and subcortical regions (striatum, pallidum, and hypothalamus). In addition, we observed increased correlations in the brain regions canonically involved in the fear memory networks, including right amygdala and right hippocampus. In contrast, the left amygdala and bilateral prefrontal cortex showed reduced functional connectivity to other brain regions, while functional connectivity of left hippocampus was not significantly different. We anticipated that *Mov10* Deletion mice would show enhanced connectivity from canonical fear centers to the periaqueductal gray area midbrain regions important for mediated the fear response (19). Interestingly, the significantly different functional connectivity of the canonical fear network regions did not include the midbrain. In contrast midbrain was overconnected to multiple cortical regions (Fig. 3D). Thus, our data suggest that the increased freezing of *Mov10* Deletion mice is stimulated by the canonical fear network directly via activation of cortical areas that then mediate freezing behavior through the midbrain indirectly via cortical regions. The average functional correlation per ROI showed that multiple cortical regions, including left visual, right gustatory, right visceral, right auditory, and right orbital regions were highly connected to the midbrain (Supplementary Fig. 3G). These results suggest that the *Mov10* Deletion mice associate foot shock and context via mechanisms that involve direct connections between the cortical areas involved in information processing and storage and the midbrain, which is responsible for eliciting the fear response.

We found enhanced functional activation of the hippocampus to cortical regions in *Mov10* Deletion mice. To explore the basis of this enhanced function connectivity and associated fear memory, we focused on hippocampal changes. As noted earlier, genes encoding ion channels have been implicated in fear memory (6,40). In addition, genes encoding channel subunits are among the MOV10-dependent differentially expressed genes in P0 hippocampus (4). To examine the intrinsic electrical properties of hippocampal neurons on a single cell level we performed whole-cell patch clamp recordings and found no difference in membrane properties and firing rate between WT and *Mov10* Deletion neurons (Supplementary Fig. 4C-F). We then examined action potential properties and found no change in overall waveform (Fig. 4B and C and Supplementary Table 2); however, we did observe that the amplitude was significantly lower in the *Mov10* Deletion neurons compared to WT both at the rheobase (Fig. 4A and B) and at 120 pA stimulation (Fig. 4C and D). The amplitude of the action potential depends on multiple factors like kinetics and density of the voltage-gated sodium channels (41–44), overall maturation state of the neuron (45), and neuronal morphology (46). We have already shown that MOV10 is required for normal dendritic morphology (4). Thus, any or all of those features could be participating in the reduced amplitude observed in the *Mov10* Deletion hippocampal neurons.

**Figure 4.**
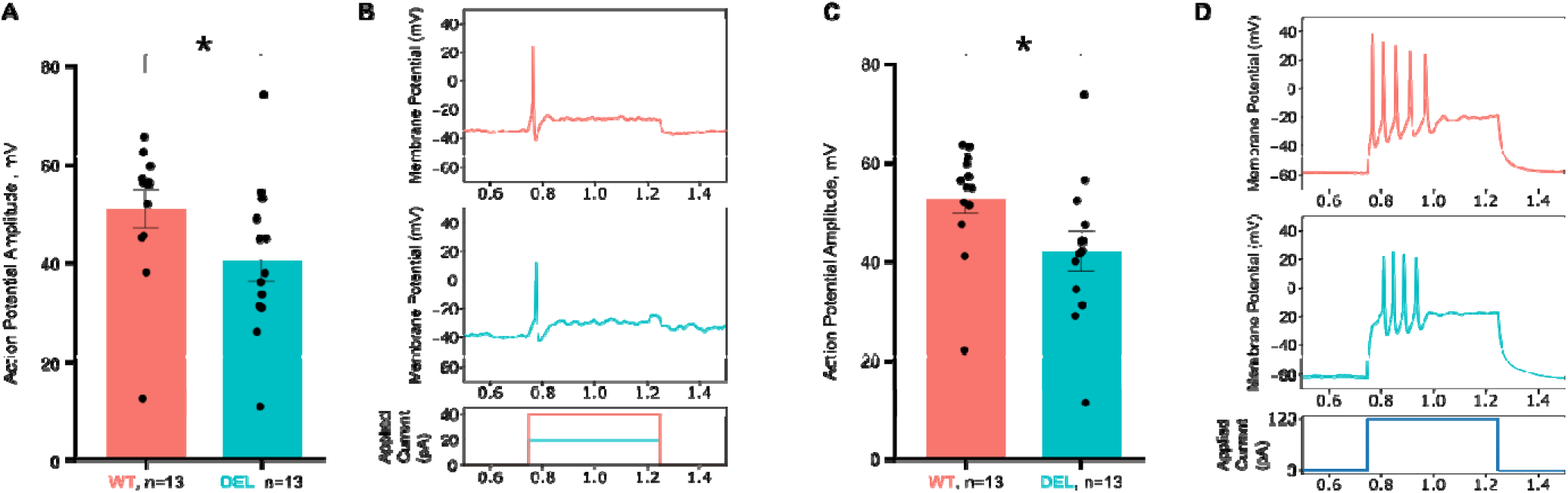
*Mov10* Deletion hippocampal neurons have reduced actional potential amplitude. **A)** Action potential amplitude of WT and *Mov10* Deletion DIV14 cultured hippocampal neurons generated by rheobase current. Y-axis is in millivolts (mV) and X-axis is genotype. **B)** Representative traces from DIV14 WT and *Mov10* Deletion neurons patch at whole-cell configuration, stimulated with rheobase current (bottom panel, picoamperes [pA]); top panels, Y axis is membrane potential (mV). **C)** Action potential amplitude (mV) of WT and *Mov10* Deletion neurons generated by 120 pA current. **D)** Representative traces from DIV14 WT and *Mov10* Deletion neurons patch at whole-cell configuration and stimulated with 120 pA current. Data are shown as mean□±□SEM. *p-value < 0.05. n is the number of neurons of the genotype indicated, both from *N*□>□3 cultures.

Since the release of neurotransmitters depends on activation of voltage-gated calcium channels, the reduced amplitude of the *Mov10* Deletion action potentials might not be high enough to trigger these channels. Thus, the consequent impaired neurotransmitter release could lead to reduced synaptic activity (47–49). To test this hypothesis, we examined the network activity of neurons cultured on microelectrode arrays (MEAs) in response to different treatments by recording activity before and after application. We expressed the drug effect as the log-transformed ratio to the baseline activity after determining that the drug effect was multiplicative rather than additive (Supplementary Fig. 5A). We began by examining the role of the glutamate receptors by treating with group 1 mGluR agonist dihydroxyphenylglycine (DHPG). While none of the treatments had a differential effect on the total number of neuronal spikes (Fig. 5A), DHPG had a profound effect on the bursting pattern, significantly increasing the number of bursts in WT compared to *Mov10* Deletion neurons (Fig. 5B); however, the number of spikes per burst and the burst duration were significantly higher in the *Mov10* Deletion neurons compared to WT (Fig. 5C and D). These effects can also be visualized in the raster plots (Fig. 5E and F). Although the number of bursts represented by red boxes appears to be higher at baseline in the *Mov10* Deletion culture, it is not significant (Supplementary Fig. 4G). However, the increased width of the red boxes in the *Mov10* Deletion culture illustrates the increased burst duration (Fig. 5E and F). These results suggest that while the absence of MOV10 from hippocampal neurons does not change the level of activity of the neurons, the pattern of activity is changed based on the type of receptor being stimulated. We tried to determine which group 1 mGluR receptor was mediating this effect using inhibitors, but the results were inconclusive (Supplementary Fig. 5D), suggesting that simultaneous stimulation of both mGluR1 and mGluR5 is required to change the activity pattern of the *Mov10* Deletion hippocampal neurons. DHPG treatment also increased the synchrony of the neuronal firing in WT neurons compared to *Mov10* Deletion neurons (Fig. 5G).

**Figure 5.**
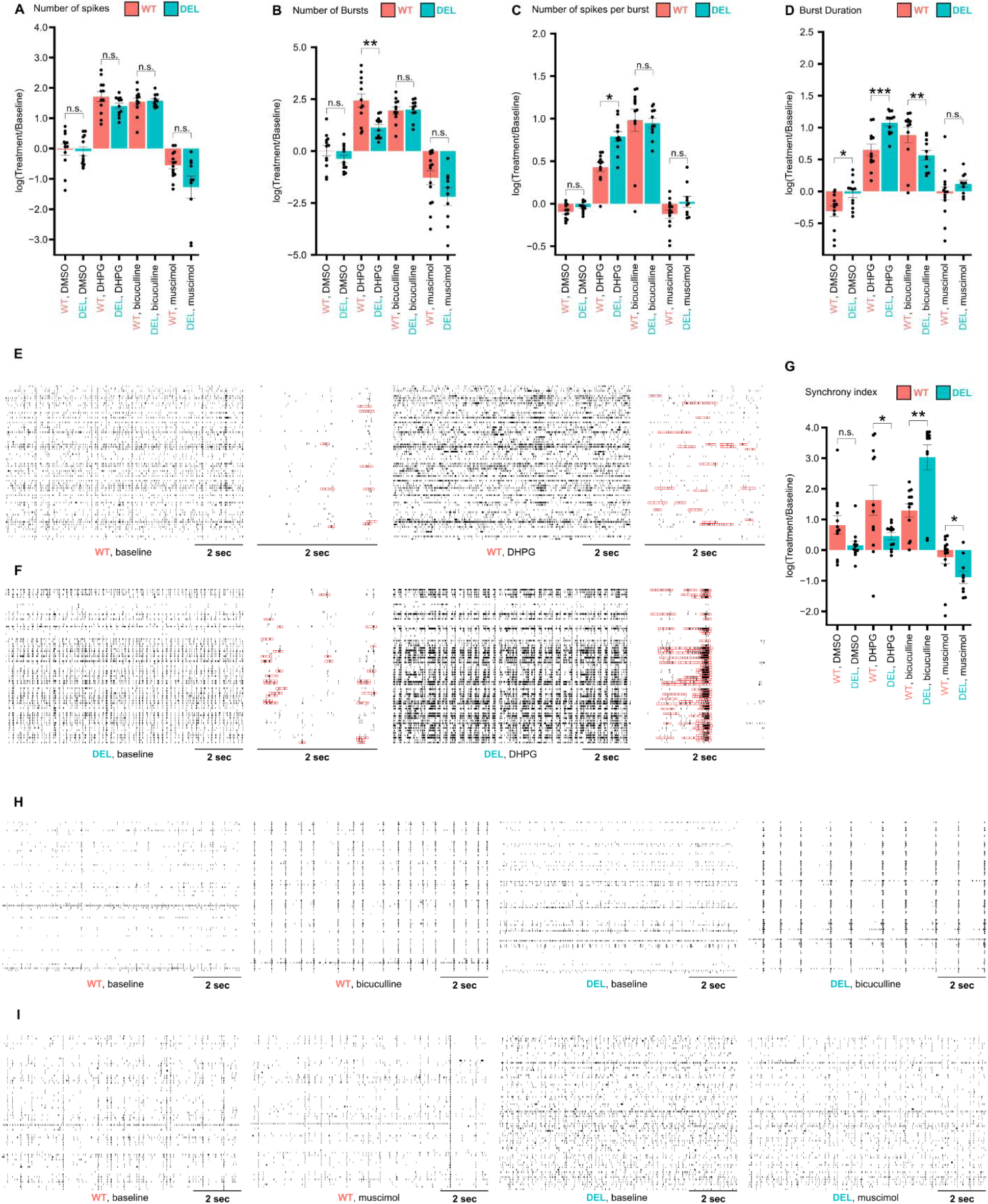
*Mov10* Deletion neurons have an altered bursting pattern after group 1 mGluR stimulation and increased synchrony after blocking inhibitory receptors. **A)** Y-axis-log fold change in spike number between Baseline and Treatment for DIV14 WT and *Mov10* Deletion hippocampal neurons measured across the 15 minutes of recording. X-axis-treatments, DMSO-solvent, DHPG-group 1 mGluR inhibitor; bicuculine-antagonist of GABA receptors; muscimol-agonist of GABA receptors. **B)** Y-axis-log fold change of burst number between Baseline and Treatment for DIV14 WT and *Mov10* Deletion hippocampal neurons measured across the 15 minutes of recording. **C)** Y axis-log fold change of spike number per burst between Baseline and Treatment for DIV14 WT and *Mov10* Deletion hippocampal neurons measured across the 15 minutes of recording. **D)** Y axis-log fold change of burst duration between Baseline and Treatment for DIV14 WT and *Mov10* Deletion hippocampal neurons measured across the 15 minutes of recording. **E-F)** Representative raster plots of the last 10 seconds of spiking activity of DIV14 WT (E) and *Mov10* Deletion (F, DEL) hippocampal neurons before and after DHPG treatment. Every vertical dash represents a single spike. Red boxes represent bursting events indicating the increased number of short bursts of WT hippocampal neurons compared to a smaller number or long duration bursts in DEL. **G)** Log fold change of synchrony index between Baseline and Treatment for DIV14 WT and *Mov10* Deletion (DEL) hippocampal neurons measured across the whole 15 minutes of recording. **H)** Representative raster plots of the last 10 seconds of spiking activity of DIV14 WT and *Mov10* Deletion hippocampal neurons before and after bicuculline treatment. Every dash represents a single spike. **I)** Representative raster plots of the last 10 seconds of spiking activity of DIV14 WT and *Mov10* Deletion hippocampal neurons before and after muscimol treatment. Every dash represents a single spike. Data are shown as mean□±□SEM from n>4 neuronal cultures from N>3 biological replicates. *p-value<0.05, **p-value<0.01.

We next turned our attention to GABA. Bicuculline, the antagonist of the GABA receptors, increased the activity of both WT and *Mov10* Deletion neurons; however, the effect on synchrony was significantly more pronounced in the *Mov10* Deletion neurons (Fig. 5G). In line with this observation, the GABA receptor agonist muscimol had the opposite effect—it reduced the synchrony in both WT and *Mov10* Deletion neurons, but significantly more for the *Mov10* Deletion neurons (Fig. 5G). This effect can also be seen in the representative raster plots (Fig. 5H and I). Upon bicuculine treatment, both WT and *Mov10* Deletion neurons fired more synchronously as indicated by an increased number of the vertical lines comprised of the aligned dashes each representing a spiking event. However, the baseline level of synchrony activity was lower in the *Mov10* Deletion culture than WT (Fig. 5H, lower number of vertical lines). Thus, the change caused by the inhibition of GABA neurons was more pronounced in the *Mov10* Deletion neuron culture (Fig. 5H). Similar to Fig. 5G, the raster plots of the cultures treated with muscimol showed the opposite effect (Fig. 5I) with the *Mov10* Deletion culture showing more of a decrease than WT. In conclusion, the *Mov10* Deletion hippocampal cultures are more responsive to inhibitory signals than WT and show a difference in the bursting activity upon stimulation of the group 1 mGluRs.

In a previous study, we found 4,054 differentially expressed genes in P0 hippocampi isolated from *Mov10* Deletion and WT mice (4). To expand this study to additional timepoints, we performed RNA seq on hippocampi isolated at P0, P7, P14, and P23. The principal component analysis showed the largest portion of the variance is time point and the second is technical replicate, suggesting that genotype explains less than 5% of the variance (Supplementary Fig. 5G). In line with this observation, the overall pattern of gene expression across different timepoints was similar in both *Mov10* Deletion and WT samples revealing three distinct clusters with characteristic expression patterns: 1) genes having high expression early and low expression later; 2) genes expressed at the same level throughout; 3) genes having low expression early and high expression later (Fig. 6A). The gene ontology analysis showed that cluster 1 has overrepresented categories that pertain to DNA-related processes, cluster 2 has overrepresented categories that pertain to transcription, and cluster 3 has overrepresented categories that describe processes important for neuronal maturation (Supplementary Fig. 5H-J). This gene expression pattern suggests that early in development neurons undergo chromatin remodeling, which drives the final transcriptional landscape. Later in development, the genes required for proper neuronal differentiation, arborization, and synaptogenesis are expressed while the housekeeping genes are expressed continuously.

**Figure 6.**
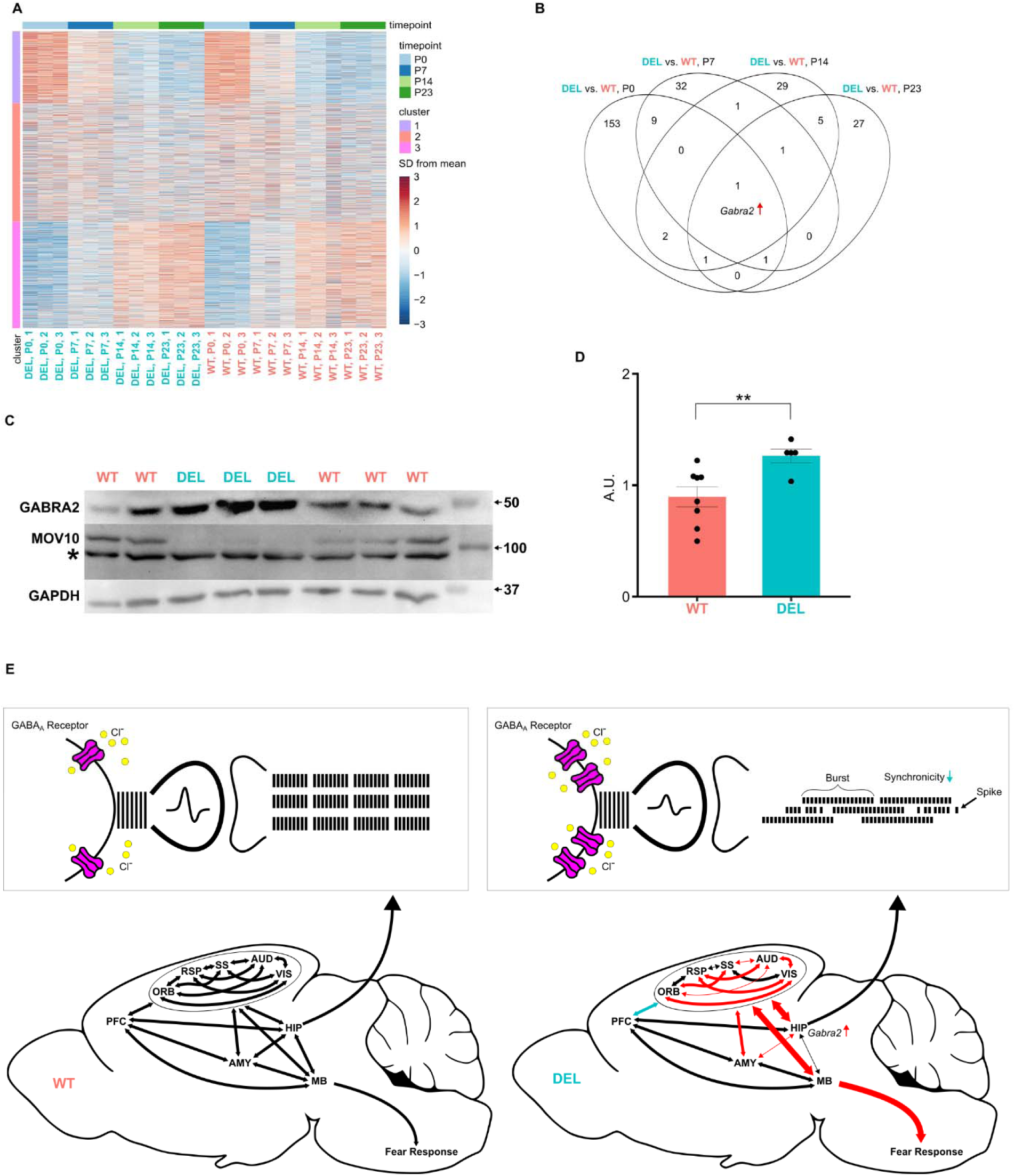
*Gabra2* expression is increased in the *Mov10* Deletion hippocampus, impacting its role in the fear network. **A)** Clustering of genes by expression pattern across time in WT and *Mov10* Deletion. **B)** Venn diagram of differentially expressed genes between WT and *Mov10* Deletion at each timepoint. **C-D)** Immunoblots of P1 hippocampal lysates probed for GABRA2, MOV10, and GAPDH; quantification of GABRA2 expression normalized to GAPDH. Asterisk indicates a non-specific band. Data are shown as mean□±□SEM from n>4 biological replicates from N>2 litters. **p-value<0.01. **E)** Unified model of WT and *Mov10* Deletion (DEL) brain activity. *Top.* Cartoon of hippocampal synapse of WT and *Mov10* Deletion showing GABAA receptor expression and the amplitude of the action potential produced. The post-synapse shows the nature of the burst pattern. The dashes represent the spiking activity of the hippocampal neurons, with *Mov10* Deletion neurons having longer bursts, smaller action potential amplitudes, and reduced synchrony. *Bottom.* Cartoon of brain connectivity of WT and *Mov10* Deletion: Cortical areas, hippocampus and amygdala have increased functional connectivity (red arrows), reduced structural connectivity (thin arrows), leading to increased stimulation of MB with consequent increased freezing behavior. Curved arrows to insets indicate the distinct bursting patterns produced by WT and *Mov10* Deletion hippocampal neurons. PFC-Prefrontal Cortex; ORB-orbital cortex; SS-Somatosensory cortex; AUD-Auditory cortex; VIS-visual cortex; RSP-retrosplenial cortex; HIP-hippocampus; AMY-Amygdala; MB-Midbrain.

Over the time course, 266 genes were differentially expressed (DE) (Fig. 6B). The only gene ontology category that was overrepresented in the DE genes was transmembrane transporter activity (Supplementary Fig. 6K). This category included genes that determine the electrical properties of the neuron (45–48) and it is possible that their concurrent dysregulation led to the reduced action potential amplitude and changed firing activity of *Mov10* Deletion neurons (Fig. 4 and 5). Importantly, one gene in the transmembrane transporter category, *Gabra2*, was the only significantly upregulated gene in the *Mov10* Deletion hippocampi at all timepoints (Fig. 6B). The *Gabra2* gene encodes the alpha-2 subunit (GABRA2) of the pentameric GABA_A_ receptor, a ligand-gated chloride channel that mediates fast inhibitory neurotransmission. Examination of the GABRA2 levels in P2 hippocampal lysates supported this result, showing significantly elevated GABRA2 levels in the *Mov10* Deletion hippocampi at P1 (Fig. 6C and D, Supplementary Fig. 5L). Therefore, our results suggest that the absence of MOV10 causes a noticeable shift in the hippocampal transcriptome over time, including increased expression of a subunit of the inhibitory GABA_A_ receptor, in part explaining the MEA results.

In summary, the *Mov10* Deletion mouse has enhanced fear memory learning, manifested by increased freezing and associated with increased functional connectivity in the brain. The canonical fear network, specifically the hippocampus and amygdala, was involved in processing fear memory, and midbrain that is responsible for eliciting freezing behavior was overconnected to cortical regions. The aberrant involvement of the hippocampus can be explained at least partially by changed electrical properties of the neurons and increased expression of *Gabra2*.

## DISCUSSION

We have identified a novel role for MOV10 in neuronal development that impacts the structural and functional connectivity in the adult mouse brain. The only condition where this novel adaptation was revealed is in the highly aversive stimulus of foot shock, which stimulates many more brain areas than do mild stressors like new environment (51). The result is that the *Mov10* Deletion mouse has a stronger associative memory for the context. It is important to note that the *Mov10* Deletion mouse does not have increased anxiety or hyperactivity (4).

The primary role of the hippocampus in fear memory formation is to integrate spatial information with aversive stimuli such as foot shock (52). However, in *Mov10* Deletion mice, this function is disrupted due to overall lower structural connectivity. This impairment manifests as an accelerated learning rate that could be associated with pathological fear expression (15–17). During fear acquisition, *Mov10* Deletion mice form an exaggerated association between the environment and the foot shock, rapidly concluding that the training chamber is dangerous. It is worth reiterating that the enhanced learning ability of the hippocampus in the absence of MOV10 manifested only after the intense stimulation of foot shock in fear conditioning (51) and remained conspicuously absent during the subtler cognitive demands of spatial exploration in Y-and T-mazes or the innate investigative behavior assessed in the novel object recognition paradigm (4).

Increased functional connectivity of hippocampus and amygdala after fear memory retrieval was observed (Fig. 3D) and expected given the evidence of their involvement in lesion studies (53,54) and the presence of engram neurons in these regions (20,55). However, the overactivation of cortical regions was unexpected and suggests that in *Mov10* Deletion mice, the fear memory, once formed, remains pathologically persistent in the cortex (56–59). Consequently, memory retrieval triggers heightened cortical activation that overstimulates the midbrain through direct inputs (60) and drives an increase in freezing behavior (Fig. 6E, red arrows). It remains unclear whether the observed differences in functional connectivity emerge immediately following fear conditioning or only after fear memory reactivation. Future studies will explore fMRI on fear-conditioned mice without memory retrieval to identify the time course of this augmentation. In addition, fear acquisition was shown to modulate sensory input, so it is possible that *Mov10* Deletion mice have increased sensory weighting (61–63). The lack of modulation from the PFC (64–67), which showed decreased functional connectivity with the cortical regions (Fig. 6E, blue arrow), further exacerbates this process.

The role of the hippocampus, which is included among the brain regions with significantly increased correlations (R HIP), might be explained by the electrophysiological experiments (Fig. 5). Release of neurotransmitters is tightly linked to the amplitude of the action potentials (47–49), suggesting that the reduced action potential amplitude in *Mov10* Deletion neurons could lead to inefficient release of synaptic vesicles. However, the results of the MEA recordings show that *Mov10* Deletion neurons can display longer bursting activity than WT neurons, supporting their ability to be stimulated (Fig. 5D). MEA electrodes typically record activity from multiple neurons, meaning that MEA bursts reflect repeated stimulation of several neurons rather than the continuous spiking of a single neuron. Increased duration of the MEA bursts suggest that *Mov10* Deletion neurons can sustain longer periods of repeated stimulation, suggesting that synaptic release in *Mov10* Deletion neurons is not impaired, despite the decreased amplitude. This could be because the decreased action potential amplitude was sufficient to trigger calcium channels, or calcium influx was adjusted to support normal synaptic release (47,68). The increased duration of the bursts suggests that *Mov10* Deletion neurons are able to activate multiple synaptic transmissions over a longer time, perhaps due to an increased pool of the synaptic vesicles (69).

At the molecular level*, Mov10* Deletion hippocampal neurons express higher levels of inhibitory GABRA2 (Fig. 6C and D). Further, stimulation with muscimol resulted in decreased synchrony of neuronal activity in the MEA experiments (Fig 5G-I), likely due to dysregulated spike timing, which is one of the major roles of inhibitory inputs (70). Our results corroborate a previous finding that GABRA2 controls network oscillations (71). Thus, the decreased synchronicity upon stimulation of the α2-containing GABA-A receptors in the absence of MOV10 might aid pattern separation (72), a computational process by which the hippocampus transforms overlapping inputs into distinct representations (72–74), resulting in the *Mov10* Deletion mice having stronger integration of contextual information during fear memory formation. Enhanced fear memory can also be supported by GABA_A_ receptors through facilitating LTP in the hippocampus (26).

Results of previous studies have found dysregulated GABRA2 in other mouse models with enhanced fear memory (75,76). The expression of genes encoding the GABA_A_R subunits can be altered at multiple levels including transcription, splicing, mRNA stability, translation, and degradation (77). Because MOV10 is an AGO2 cofactor, we suspect that MOV10 facilitates AGO2 association with the 3’UTR of *Gabra2* leading to translation suppression and degradation. Thus, in the absence of MOV10, as in the *Mov10* Deletion hippocampus, *Gabra2* expression is increased because AGO2 is unable to access its 3’UTR. Alternatively, it is also possible that MOV10 may indirectly regulate GABRA2 expression by directly regulating a protein that regulates *Gabra2* expression or stability (78).

The structural connectivity in the brain reflects the integrity of white matter structures like axonal tracts. Axonal development in the *Mov10* Deletion mouse would be predicted to be impaired by our previous work showing that MOV10 regulates expression of the microtubule binding protein NUMA1 by modulating AGO2 access to its 3’UTR (4). NUMA1 has been shown by others to be required for normal axonal projections in the corpus callosum, functioning in the axonal growth cone (79). Thus, the decrease in structural connectivity of cortical regions in the absence of MOV10 might initiate compensatory mechanisms that ultimately result in the increased functional connectivity, similar to what has been observed in models of multiple sclerosis and traumatic brain disorder (80,81) that manifest upon the strong sensory stimulation of foot shock.

The implications of this work for human mental health are significant. Recently, MOV10 was identified in an epigenome-wide study of individuals with substance dependence (82). It is tempting to speculate that individuals with reduced MOV10 expression may experience enhanced fear memory, which leads to unhealthy coping strategies like tobacco, alcohol or cocaine use. Interestingly, GABRA2 was shown to be linked to alcohol dependence both though genomic and electrophysiological studies (83,84). In addition, a Genome Wide Association Study (GWAS) of a million veterans with PTSD identified a number of genes including *CAPZA1* (85) as co-occurring in patients. CAPZA1 is less than 5 kilobases from MOV10 (86), raising the intriguing possibility that expression of the linked MOV10 gene participates in PTSD risk. Thus, identification of the neuronal networks involved in augmented fear memory will provide a deeper understanding of the underlying causes of substance dependence, PTSD and autism, in which enhanced fear memory impacts the quality of life (87–89).

## ACKNOWLEGEMENTS

We would like to acknowledge Shreyan Majumdar, Brad Sutton, Paul Camacho, Tracey Wszalek, Biomedical Imaging Center at Beckman Institute and Zhiyu Kuang for help with MRI data acquisition and analysis, Dr. Yeeun Yook for the help with MEA data acquisition and analysis, Dr. Dave Zhang, Rachan Raj, Ivy Yang, and Biostate.ai for assistance with the RNA-sequencing.

## DATA AVAILABILITY

All data RNA-sequencing data is available at GEO database through accession number GSE317128. All the other data generated in this study are available from the corresponding author upon request.

## Supplementary Materials

**Supplementary Figure 1.**
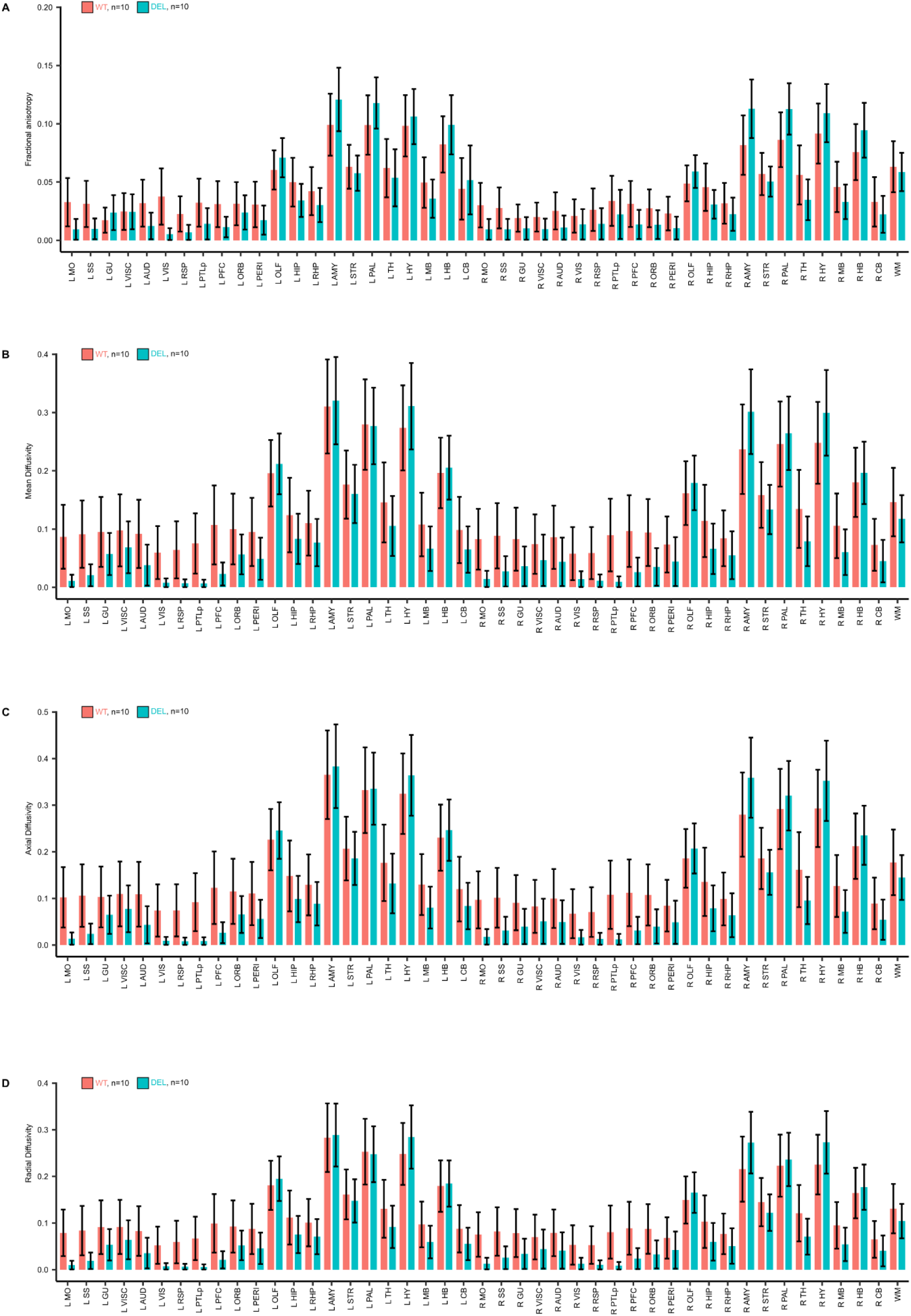
Diffusion tensor metrics are not significantly different in *Mov10* Deletion mice. **A)** Average fractional anisotropy from n=10 DTI scans of WT and *Mov10* Deletion mice across 22 bilateral regions and white matter (WM). **B)** Average mean diffusivity from n=10 DTI scans of WT and *Mov10* Deletion mice across 22 bilateral regions and white matter (WM). **C)** Average axial diffusivity from n=10 DTI scans of WT and *Mov10* Deletion mice across 22 bilateral regions and white matter (WM). **D)** Average radial diffusivity from n=10 DTI scans of WT and *Mov10* Deletion mice across 22 bilateral regions and white matter (WM). Data are shown as mean□±□SEM, n is the number of mice of the genotype indicated, of both sexes from *N*□>□3 litters.

**Supplementary Figure 2.**
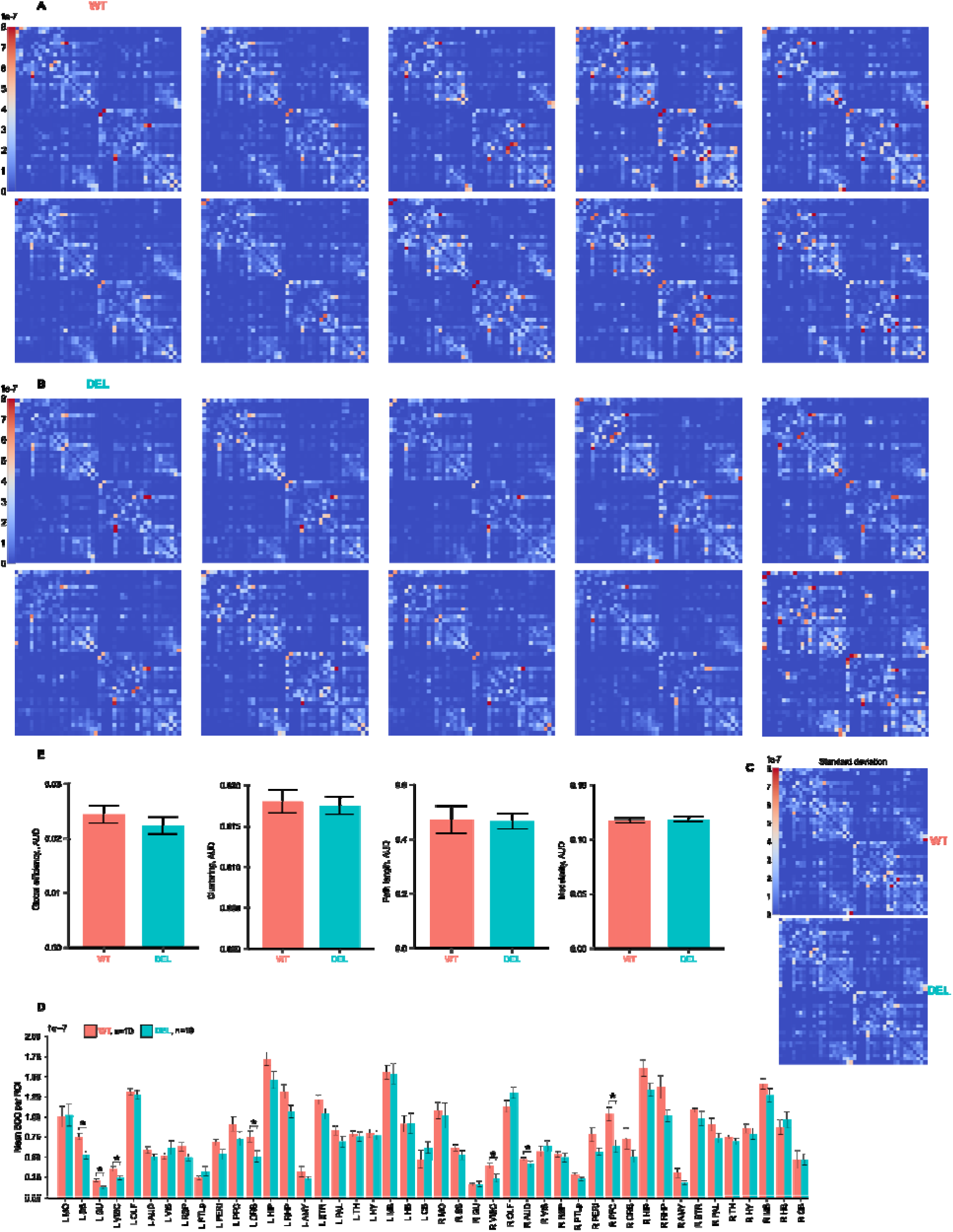
Structural connectivity and graph analysis measurements of structural networks. **A)** Connectivity matrices with strength of connectivity (SOC) between 22 bilateral brain regions from n=10 DTI scans of WT mice. **B)** Connectivity matrices with SOC between 22 bilateral brain regions from n=10 DTI scans of *Mov10* Deletion mice. **C)** Standard deviation matrices of SOC between 22 bilateral brain regions across 10 animals for each genotype. **D)** Mean SOC measures for all regions averaged across the X-axis of the connectivity matrix. **E)** Area under the curve (AUC) of graph analysis measurements calculated on the connectivity matrices with SOC across several thresholds. Data are shown as mean□±□SEM, *p-value<0.05, n is the number of mice of the genotype indicated, of both sexes from *N*□>□3 litters.

**Supplementary Figure 3.**
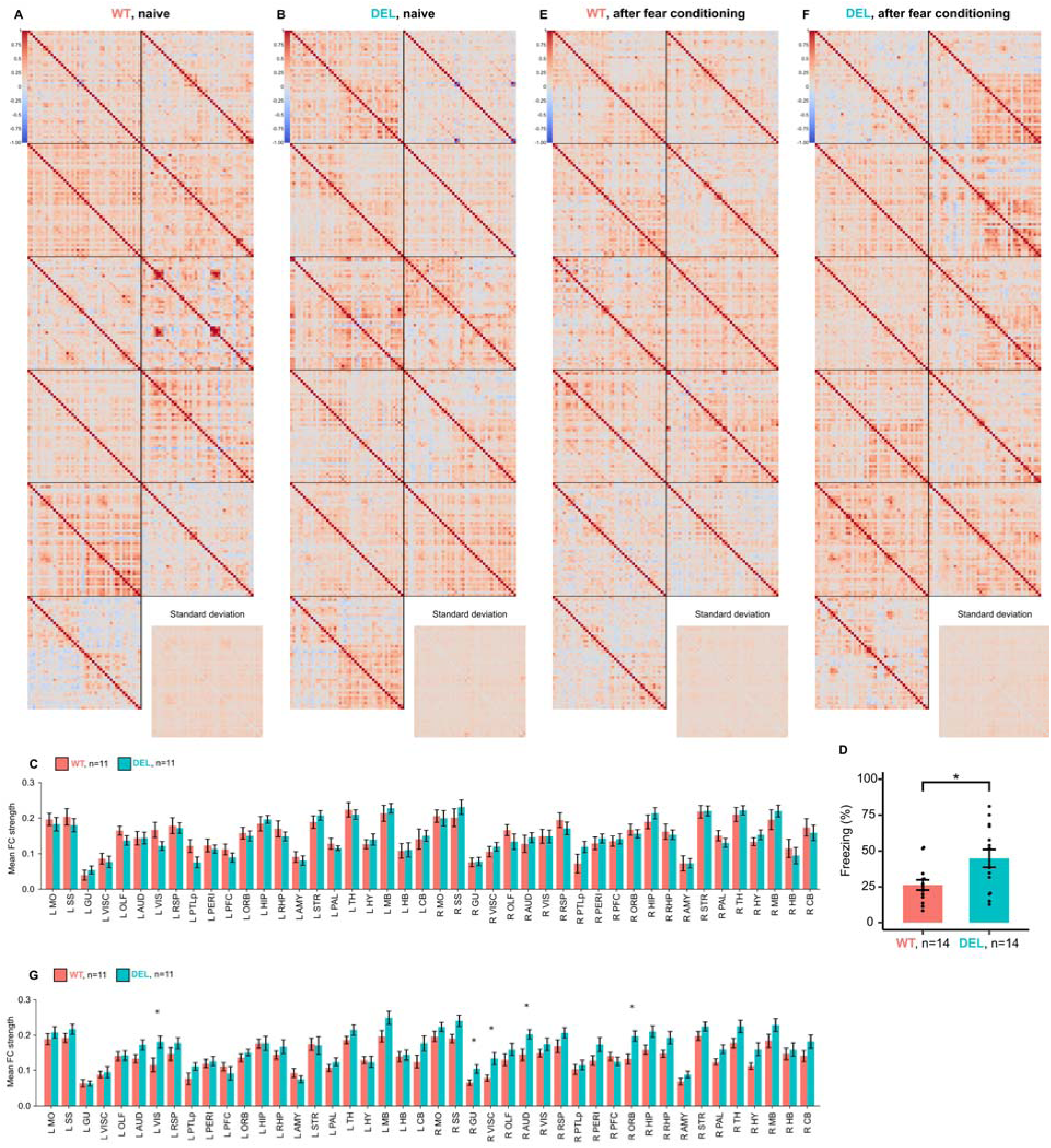
Functional connectivity of WT and *Mov10* Deletion mice. **A)** Functional connectivity and standard deviation matrices with Z-transformed Pearson’s correlation coefficients between 22 bilateral brain regions from 14 fMRI scans of naïve WT mice. **B)** Functional connectivity and standard deviation matrices with Z-transformed Pearson’s correlation coefficients between 22 bilateral brain regions from 14 fMRI scans of naïve *Mov10* Deletion mice. **C)** Mean functional correlation coefficients calculated from rs-fMRI data acquired before fear conditioning averaged across the X-axis of the connectivity matrix. **D)** Results of the context fear conditioning test on mice that were scanned using MRI. **E)** Functional connectivity and standard deviation matrices with Z-transformed Pearson’s correlation coefficients between 22 bilateral brain regions from 14 fMRI scans of WT mice after fear conditioning test. **F)** Mean functional connectivity and standard deviation matrices with Z-transformed Pearson’s correlation coefficients between 22 bilateral brain regions from 14 fMRI scans of *Mov10* Deletion mice after fear conditioning. **G)** Mean functional correlation coefficients calculated from fMRI data acquired before fear conditioning averaged across the X-axis of the connectivity matrix. Data are shown as mean□±□SEM, *p-value<0.05, n is the number of mice of the genotype indicated, of both sexes from *N*□>□3 litters.

**Supplementary Figure 4.**
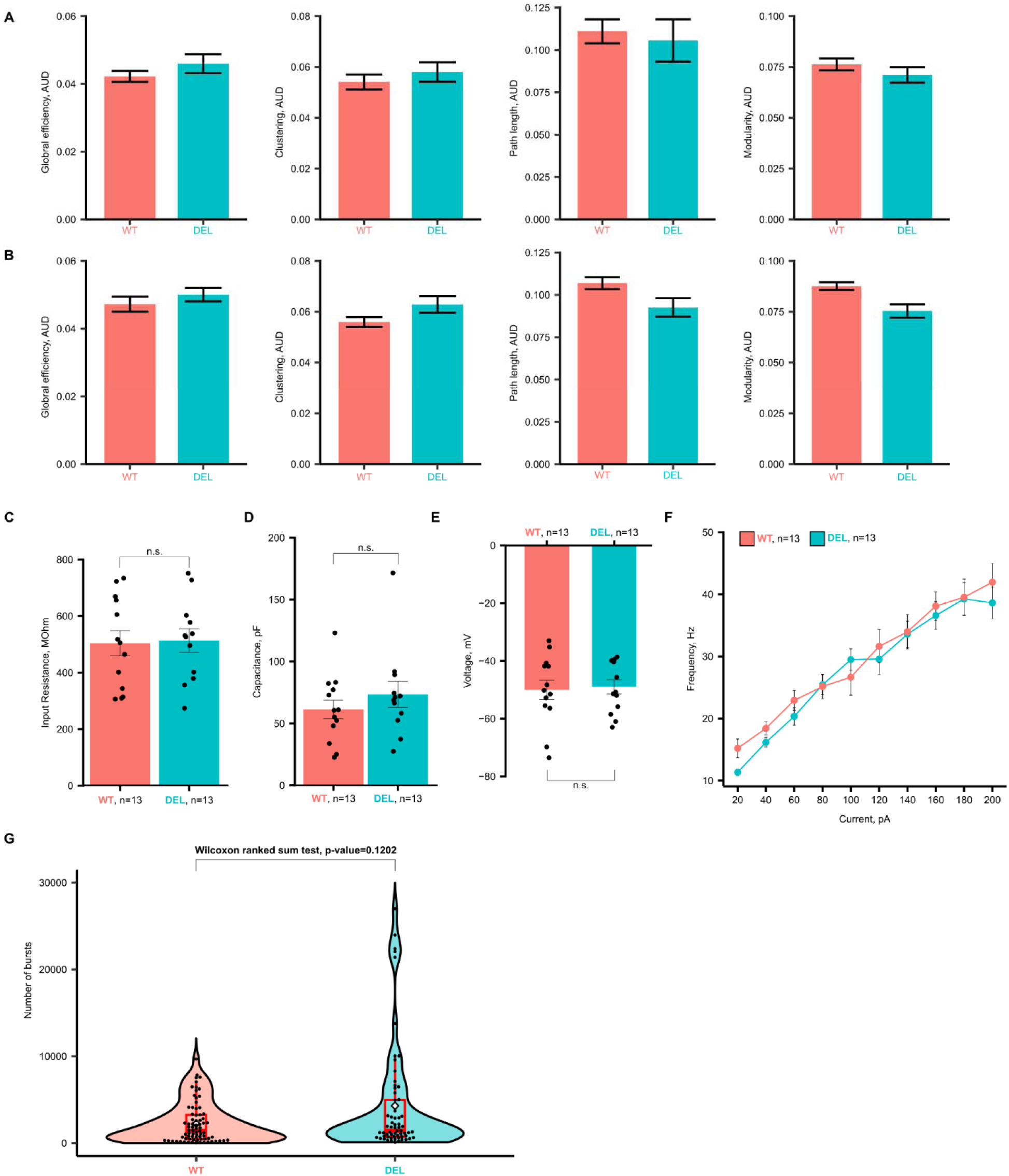
Graph analysis metrics of functional networks and number of bursts in baseline activity of hippocampal cultures are not different in *Mov10* Deletion mice. **A)** Area under the curve (AUC) of graph analysis measurements calculated on the connectivity matrices with Z-transformed Pearson’s correlation coefficients from fMRI data of naïve mice across several thresholds. **B)** Area under the curve (AUC) of graph analysis metrics calculated on the connectivity matrices with Z-transformed Pearson’s correlation coefficients from fMRI data of fear conditioned mice across several thresholds. **C)** Membrane input resistance of WT and *Mov10* Deletion (DEL) DIV14 cultured hippocampal neurons. **D)** Membrane capacitance of WT and *Mov10* Deletion DIV14 cultured hippocampal neurons. **E)** Membrane resting potential of WT and *Mov10* Deletion DIV14 cultured hippocampal neurons. **F)** Instantaneous action potential firing rate for WT and *Mov10* Deletion DIV14 cultured hippocampal neurons. Data are shown as mean□±□SEM. n is the number of neurons of the genotype indicated, both from *N*□>□3 cultures. **G)** Number of bursts in baseline recordings of WT and *Mov10* Deletion mice.

**Supplementary Figure 5.**
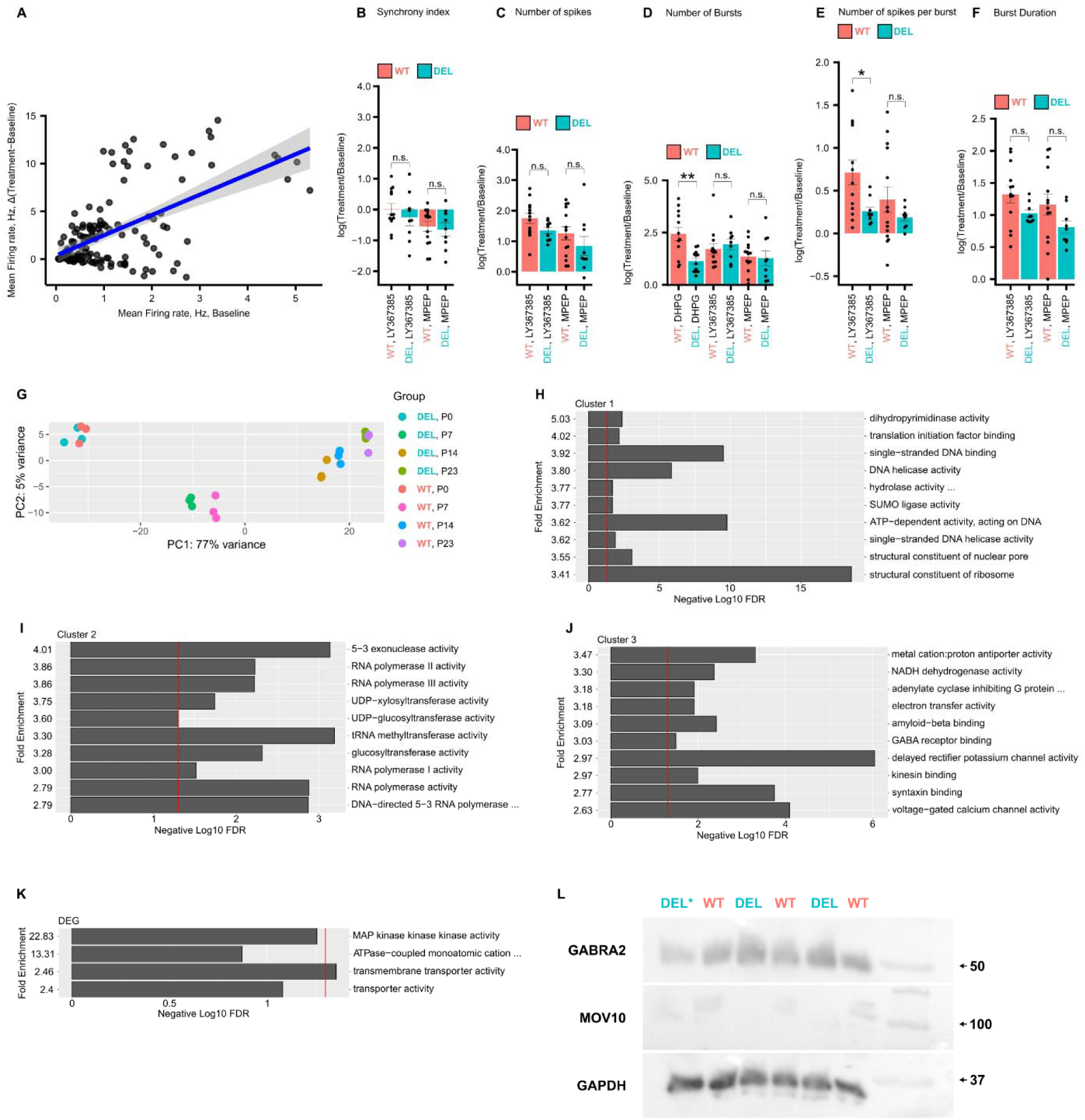
Effects of mGluR inhibitors on neuronal firing and results of gene ontology enrichment analysis. **A)** Linear regression of Mean Firing Rate between Δ(Treatment−Baseline) and Baseline. Blue line is the linear fit and shaded area around the line is the standard error. **B)** Log fold change of synchrony index between Baseline and Treatment for DIV14 WT and *Mov10* Deletion (DEL) hippocampal neurons measured across the whole 15 minutes of recording. **C)** Log fold change of number of spikes between Baseline and Treatment for DIV14 WT and *Mov10* Deletion hippocampal neurons measured across the whole 15 minutes of recording. **D)** Representative raster plots of the last 10 seconds of spiking activity of DIV14 WT and *Mov10* Deletion hippocampal neurons before and after bicuculline treatment. Every dash represents a single spike. **E)** Log fold change of number of bursts between Baseline and Treatment for DIV14 WT and *Mov10* Deletion hippocampal neurons measured across the whole 15 minutes of recording. **F)** Log fold change of number of spikes per burst between Baseline and Treatment for DIV14 WT and *Mov10* Deletion hippocampal neurons measured across the whole 15 minutes of recording. **G)** Log fold change of duration of burst between Baseline and Treatment for DIV14 WT and *Mov10* Deletion hippocampal neurons measured across the whole 15 minutes of recording. Data are shown as mean□±□SEM from n>4 neuronal cultures from N>3 biological replicates. *p-value<0.05. **H-J)** Significantly overrepresented gene ontology categories in each of the identified cluster. Red line represents the -log(0.05) = 1.3. **K)** Gene ontology categories identified in the differentially expressed genes across all timepoints. Red line represents the -log(0.05) = 1.3. **L)** Immunoblots of GABRA2, MOV10, and GAPDH. Asterisk represents outlier DEL sample that was not included in quantification (Z-score>5).

**Supplementary Table 1.** Label names of the brain regions used for MRI data analysis.

| **Parental** | **Name** | **Acronym** | **ID** |
| --- | --- | --- | --- |
| Motor | Motor | MO | 500 |
| Somatosensory | Somatosensory | SS | 453 |
| Gustatory | Gustatory | GU | 1057 |
| Visceral | Visceral | VISC | 677 |
| Auditory | Auditory | AUD | 247 |
| Visual | Visual | VIS | 669 |
| Retrosplenial | Retrosplenial | RSP | 254 |
| Posterior parietal association | Posterior parietal association | PTLp | 22 |
| Prefrontal cortex | Anterior cingulate | ACA | 31 |
| Prefrontal cortex | Prelimbic | PL | 972 |
| Prefrontal cortex | Infralimbic | ILA | 44 |
| Prefrontal cortex | Frontal pole | FRP | 184 |
| Orbital | Orbital | ORB | 714 |
| Orbital | Agranular insular | AI | 95 |
| Orbital | Clastrum | CLA | 583 |
| Perirhinal | Temporal association | TEa | 541 |
| Perirhinal | Perirhinal | PERI | 922 |
| Perirhinal | Ectorhinal | ECT | 895 |
| Olfactory | Olfactory | OLF | 698 |
| Olfactory | Endopiriform | EP | 942 |
| Hippocampus | Hippocampus | HIP | 1080 |
| Retrohippocampal region | Retrohippocampal region | RHP | 822 |
| Amygdala | Lateral amygdalar nucleus | LA | 131 |
| Amygdala | Basolateral amygdalar nucleus | BLA | 295 |
| Amygdala | Basomedial amygdalar nucleus | BMA | 319 |
| Amygdala | Posterior amygdalar nucleus | PA | 780 |
| Amygdala | Striatal amygdala | sAMY | 278 |
| Striatum | Dorsal striatum | STRd | 485 |
| Striatum | Ventral striatum | STRv | 493 |
| Striatum | Lateral septal complex | LSX | 275 |
| Pallidum | Pallidum | PAL | 803 |
| Thalamus | Thalamus | TH | 549 |
| Hypothalamus | Hypothalamus | HY | 1097 |
| Midbrain | Midbrain | MB | 313 |
| Hindbrain | Hindbrain | HB | 1065 |
| Cerebellum | Cerebellum | CB | 512 |
| WMCSF | fiber tracts | fiber tracts | 1009 |
| WMCSF | Ventricular System | VS | 73 |

**Supplementary Table 2.** Action potential properties of *Mov10* Deletion and WT hippocampal neurons.

| Rheobase |  |  |  |  |  |  |  |
| --- | --- | --- | --- | --- | --- | --- | --- |
| Genotype | Latency, ms | V_thr, mV | 10-90% rise time, ms | 90-10% decay time, ms | Half width, ms | AHP, mV | Rheobas pA |
| WT | 32.35±6.97 | -<br>25.06±1.79 | 1.15±0.16 | 2.20±0.20 | 0.59±0.04 | -21.81±1.59 | 52.31±10. |
| *Mov10* Deletion | 66.55±24.92 | -<br>20.46±2.69 | 1.17±0.07 | 2.24±0.14 | 0.65±0.04 | -21.20±1.98 | 38.46±7.6 |
| p-value | 0.06 | 0.23 | 0.47 | 0.72 | 0.36 | 0.8 | 0.28 |
| 120 pA |  |  |  |  |  |  |  |
| Genotype | Latency, ms | V_thr, mV | 10-90% rise time, ms | 90-10% decay time, ms | Half width, ms | AHP, mV |  |
| WT | 12.82±3.54 | -<br>23.59±1.95 | 1.08±0.13 | 2.49±0.26 | 0.56±0.06 | -12.80±1.84 |  |
| *Mov10* Deletion | 13.38±4.20 | -<br>16.72±3.99 | 1.12±0.09 | 2.62±0.28 | 0.64±0.05 | -12.15±1.98 |  |
| p-value | 0.98 | 0.15 | 0.39 | 0.93 | 0.26 | 0.84 |  |

