## Supplementary figures and images for "Reduced MOV10 reveals novel functional cortical connections in an increased fear response"

### Supplementary Fig. 1

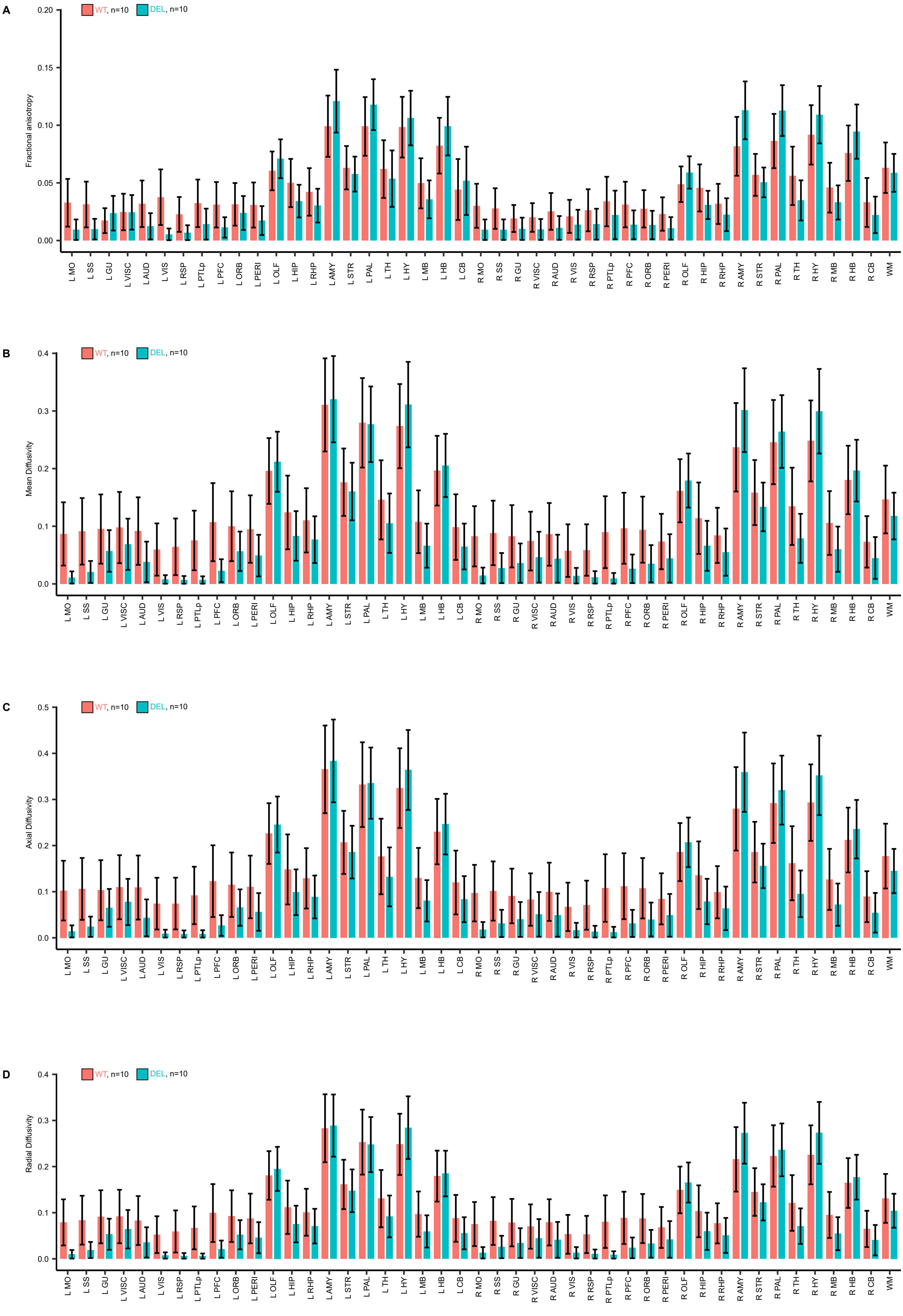

### Supplementary Fig. 2

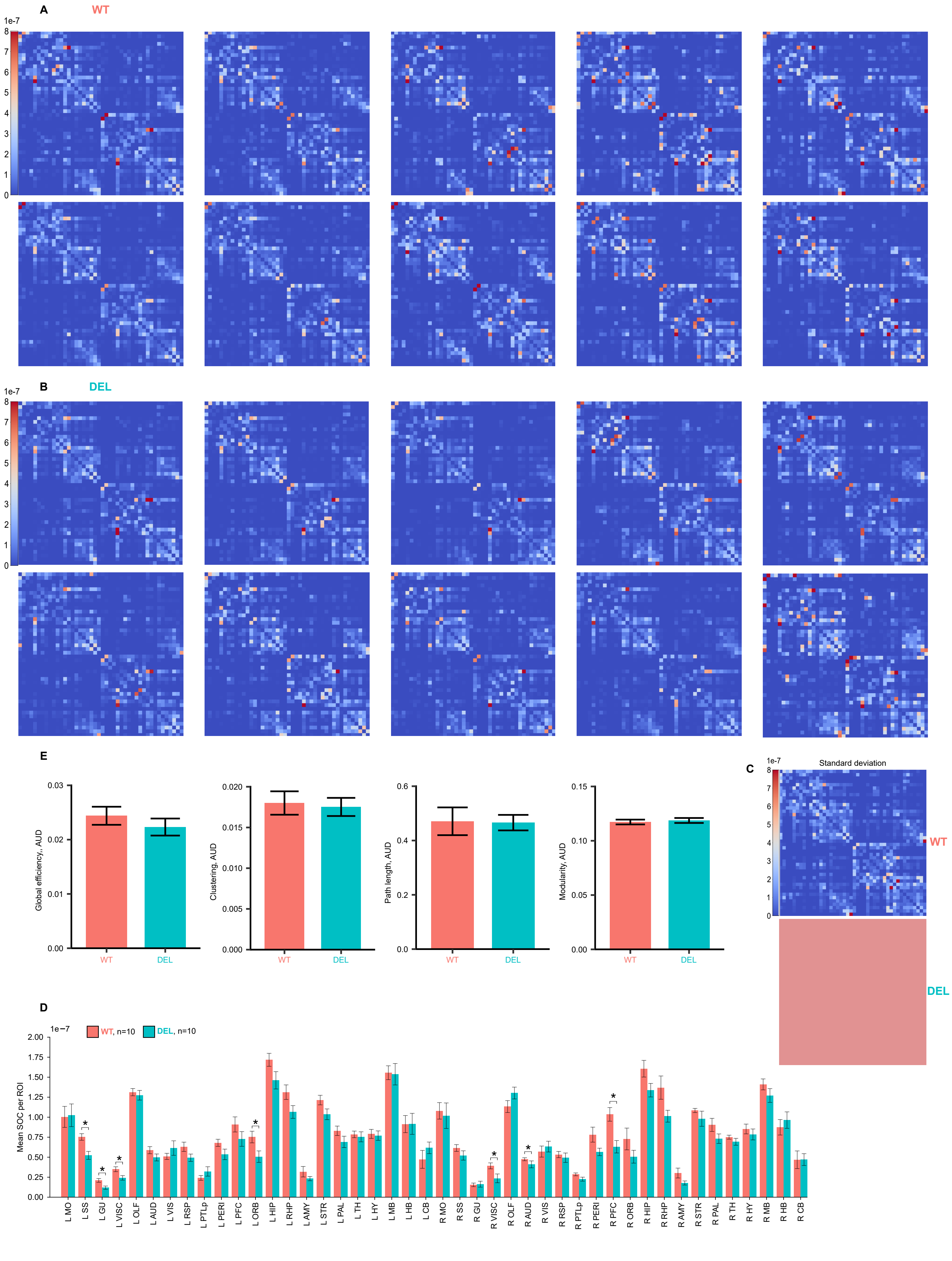

### Supplementary Fig. 3

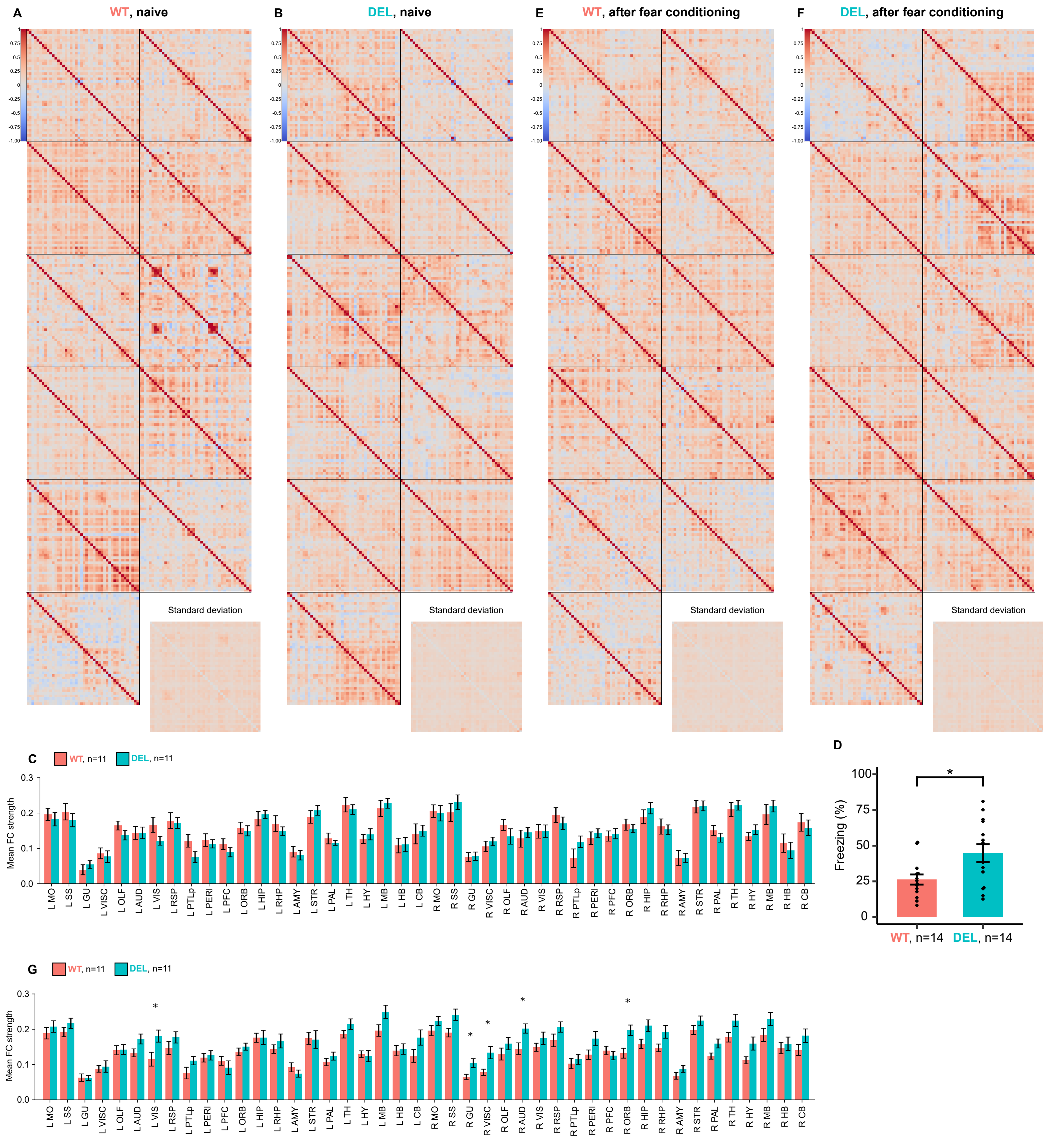

### Supplementary Fig. 4

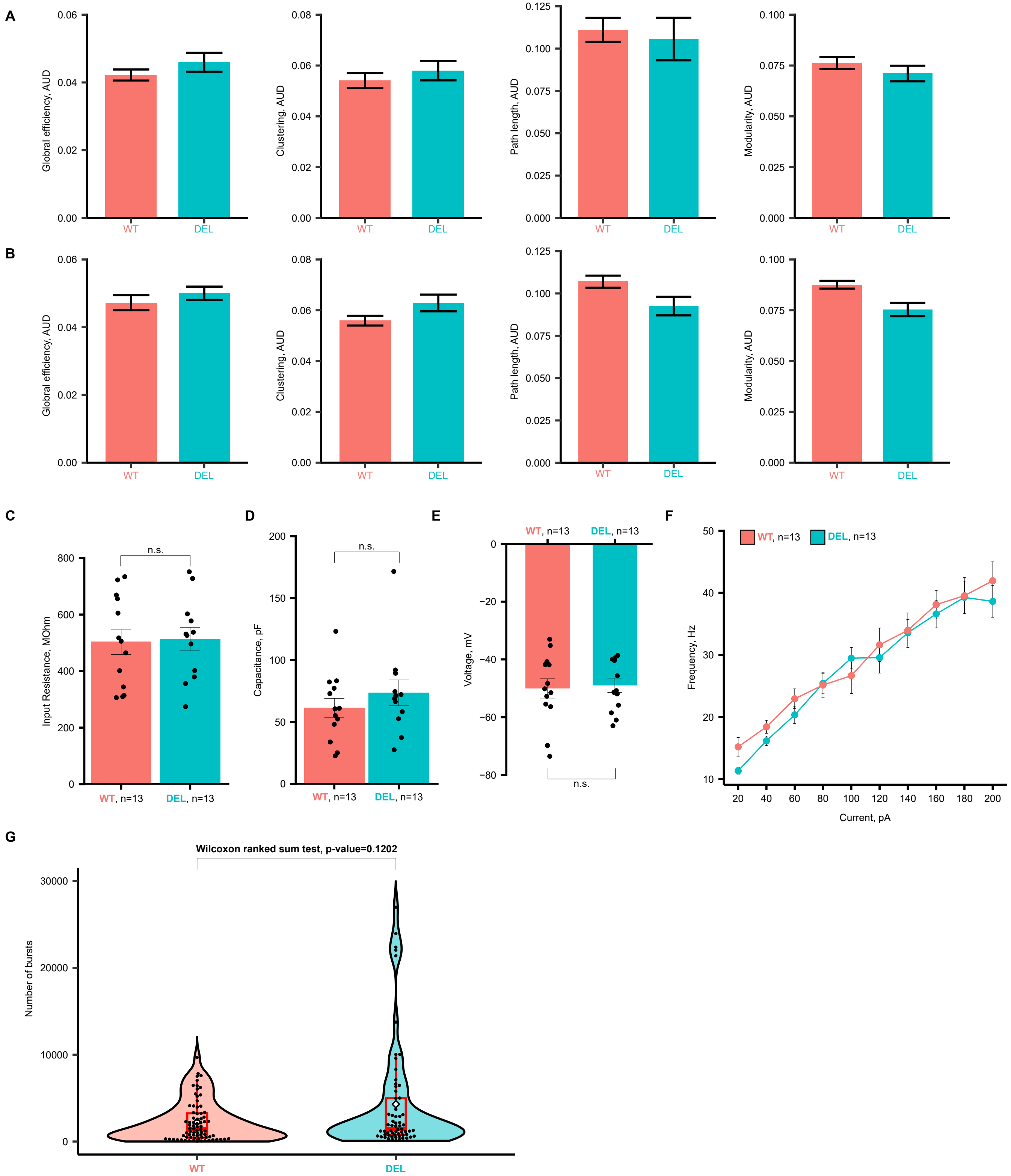

### Supplementary Fig. 5

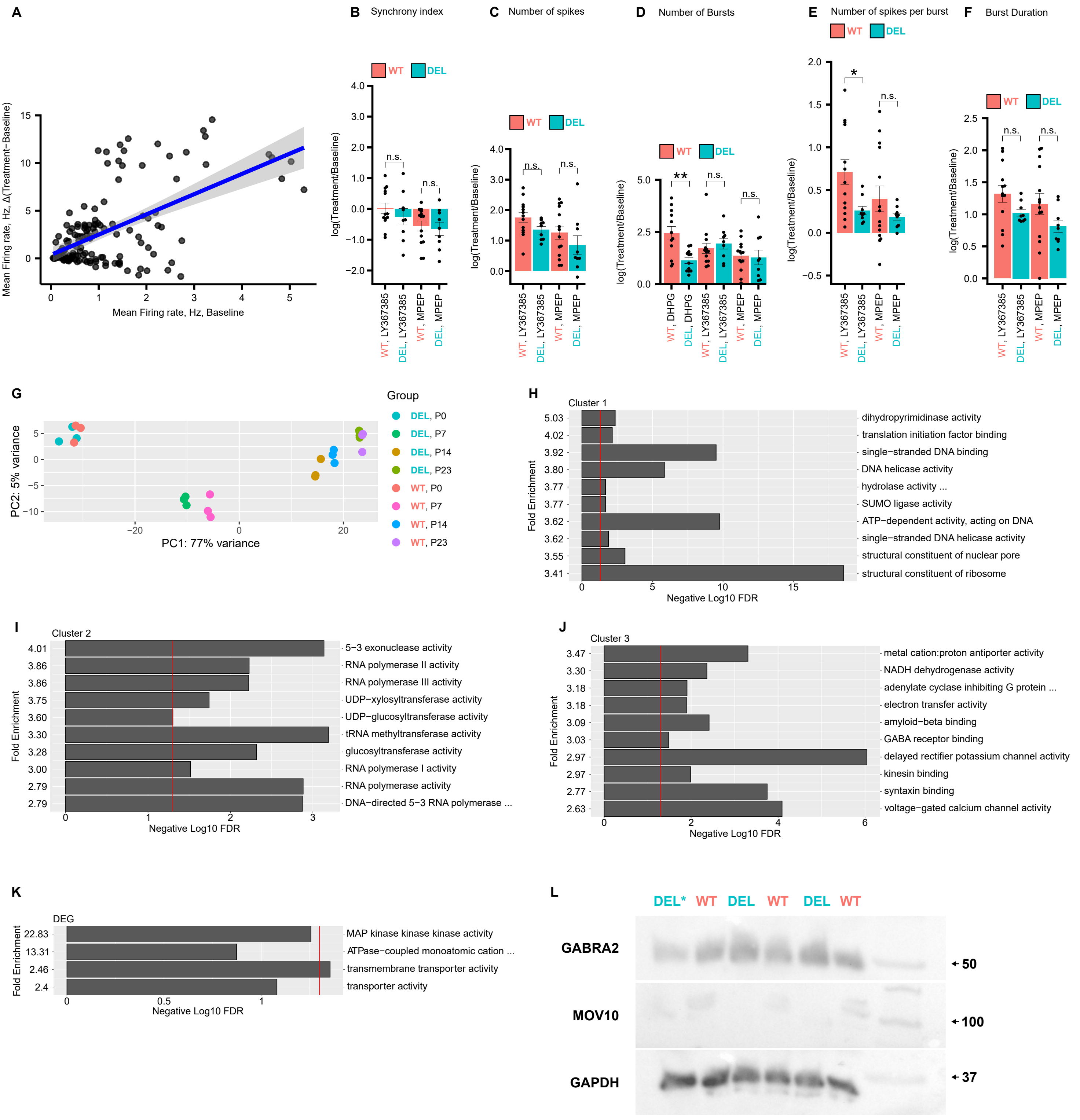
